# CLCC1 promotes hepatic neutral lipid flux and nuclear pore complex assembly

**DOI:** 10.1101/2024.06.07.597858

**Authors:** Alyssa J. Mathiowetz, Emily S. Meymand, Kirandeep K. Deol, Güneş Parlakgül, Mike Lange, Stephany P. Pang, Melissa A. Roberts, Emily F. Torres, Danielle M. Jorgens, Reena Zalpuri, Misun Kang, Casadora Boone, Yaohuan Zhang, David W. Morgens, Emily Tso, Yingjiang Zhou, Saswata Talukdar, Tim P. Levine, Gregory Ku, Ana Paula Arruda, James A. Olzmann

**Affiliations:** Department of Molecular and Cell Biology, University of California, Berkeley, Berkeley, CA 94720, USA; Department of NutriYonal Sciences and Toxicology, University of California, Berkeley, Berkeley, CA 94720, USA; Electron Microscope Laboratory, University of California, Berkeley, Berkeley, CA 94720, USA; Diabetes Center, University of California, San Francisco, San Francisco, CA 94143, USA; Merck & Co., Inc., South San Francisco, CA 94080, USA; University College London InsYtute of Ophthalmology, Bath Street London, EC1V 9EL, UK; Department of Medicine, Division of Endocrinology, University of California, San Francisco, San Francisco, CA 94143, USA; Chan Zuckerberg Biohub, San Francisco, CA 94158, USA

## Abstract

Imbalances in lipid storage and secretion lead to the accumulation of hepatocyte lipid droplets (LDs) (i.e., hepatic steatosis). Our understanding of the mechanisms that govern the channeling of hepatocyte neutral lipids towards cytosolic LDs or secreted lipoproteins remains incomplete. Here, we performed a series of CRISPR-Cas9 screens under different metabolic states to uncover mechanisms of hepatic neutral lipid flux. Clustering of chemical-genetic interactions identified CLIC-like chloride channel 1 (CLCC1) as a critical regulator of neutral lipid storage and secretion. Loss of CLCC1 resulted in the buildup of large LDs in hepatoma cells and knockout in mice caused liver steatosis. Remarkably, the LDs are in the lumen of the ER and exhibit properties of lipoproteins, indicating a profound shift in neutral lipid flux. Finally, remote homology searches identified a domain in CLCC1 that is homologous to yeast Brl1p and Brr6p, factors that promote the fusion of the inner and outer nuclear envelopes during nuclear pore complex assembly. Loss of CLCC1 lead to extensive nuclear membrane herniations, consistent with impaired nuclear pore complex assembly. Thus, we identify CLCC1 as the human Brl1p/Brr6p homolog and propose that CLCC1-mediated membrane remodeling promotes hepatic neutral lipid flux and nuclear pore complex assembly.

## Main

Lipid droplets (LDs) are the primary lipid storage organelle in cells^1,2^. LDs are derived from the endoplasmic reticulum (ER) through a process involving neutral lipid synthesis and deposition between the leaflets of the ER bilayer, neutral lipid phase separation facilitated by LD assembly complexes, and LD emergence into the cytoplasm from the outer leaflet of the ER bilayer^1,2^. Mature LDs consist of a neutral lipid core that is encircled by a phospholipid monolayer decorated with integral and peripheral proteins^1,2^. LDs act as a dynamic cellular repository of lipids that can be formed *de novo* and that can be rapidly broken down to release stored lipids^1,2^. LD degradation provides substrates for β-oxidation and the generation of energy or macromolecular building blocks for the biosynthesis of membranes and lipid signaling molecules^1,2^. LDs also suppress lipotoxicity by sequestering lipids and preventing their flux into damaging species^1,2^, such as di-saturated glycerolipids that cause ER stress^3^ or polyunsaturated fatty acid-containing phospholipids that are prone to oxidation^4,5^. LD dysregulation has been implicated in the pathogenesis of a wide variety of diseases, ranging from metabolic diseases to cancer and neurodegeneration^6–8^. In addition, the functions, composition, and regulation of LDs differ depending on the cell and tissue type, metabolic conditions, and fluctuations in the cellular need for lipids and energy^1,9^.

The liver is a central site of lipid metabolism^6,10^. Fatty acids in liver hepatocytes may be used to generate triacylglycerol (TAG) that is stored in cytoplasmic LDs or packaged into very low-density lipoproteins (VLDLs) for secretion into circulation^6,10,11^. Altered storage and secretion of lipids can lead to the persistent buildup of hepatocyte LDs, a pathological hallmark of Metabolic dysfunction-associated fatty liver disease (MAFLD)^6,12^. An estimated 20-30% of the general population and 75-92% of morbidly obese individuals exhibit hepatic steatosis^12^, and MAFLD is a risk factor for Metabolic dysfunction-associated steatohepatitis (MASH), cirrhosis, and hepatocellular carcinoma^6,12^. The mechanisms that govern the storage of neutral lipids in hepatocyte LDs remain incompletely understood, and addressing this gap in knowledge is paramount to the development of new therapeutic strategies.

In the current study, we performed a series of over 20 CRISPR-Cas9 genetic screens in Huh7 hepatoma cells under different metabolic conditions, providing a compendium of genetic modifiers of lipid storage and insights into the mechanisms that govern hepatic neutral lipid flux and cellular membrane homeostasis. Analyses of the resulting chemical-genetic interactions identify CLCC1 as a strong regulator of hepatic neutral lipid storage. We further discover that CLCC1 is the human homolog of yeast Brl1p / Brr6p, two proteins implicated in membrane remodeling and fusion during nuclear pore complex assembly^13–18^. Indeed, loss of CLCC1 results in defects in nuclear pore complex assembly and in altered neutral lipid flux, leading to hepatic steatosis due to enlarged lipoproteins that fail to be secreted and accumulate in the ER lumen at the expense of cytosolic LDs. Our results demonstrate the importance of CLCC1 for the fusion of the inner and outer nuclear envelopes during nuclear pore complex assembly and neutral lipid flux in hepatocytes.

## Results

### CRISPR-Cas9 screens identify a compendium of neutral lipid storage genetic modifiers

The fluorescent dye BODIPY 493/503 concentrates in the LD neutral lipid core, enabling quantification of changes in neutral lipids sequestered in LDs by fluorescence imaging and flow cytometry (**Fig 1A and Extended Data Fig 1A**). To systematically identify genes involved in lipid storage, we performed a genome-wide fluorescence-activated cell sorting (FACS)-based CRISPR-Cas9 screen in Huh7 hepatoma cells using BODIPY 493/503 fluorescence as a reporter of neutral lipid storage (**Fig 1B and Supplementary Table 1**). Duplicate batch retest screens were subsequently performed using a custom Validation of Lipid Droplet & Metabolism (VLDM) sgRNA library (**Extended Data Fig 1B**) to increase confidence and reduce false positives and negatives (**Fig 1C and Extended Data Fig 1C**). These data identified 244 high confidence candidate regulators of neutral lipid storage – 192 positive and 52 negative regulators. As anticipated, the screen was enriched in genes associated with glycerolipid metabolism, including factors involved in neutral lipid synthesis (e.g., *ACSL3*, *SCD*, *CHP1*, *ACACA*, *DGAT2*, *AGPAT6*) and lipolysis (e.g., *ABHD5*, *PNPLA2*, *HILPDA*) (**Fig 1C,D and Extended Data Fig 1D**), consistent with the overall quality of the screens. Genes involved in other functional categories were also identified, such as the secretory pathway and protein trafficking, ubiquitination and ERAD, the mevalonate pathway, SREBP pathway, and additional processes that influence lipid metabolism (**Fig 1C,D and Extended Data Fig 1D**).

**Figure 1.**
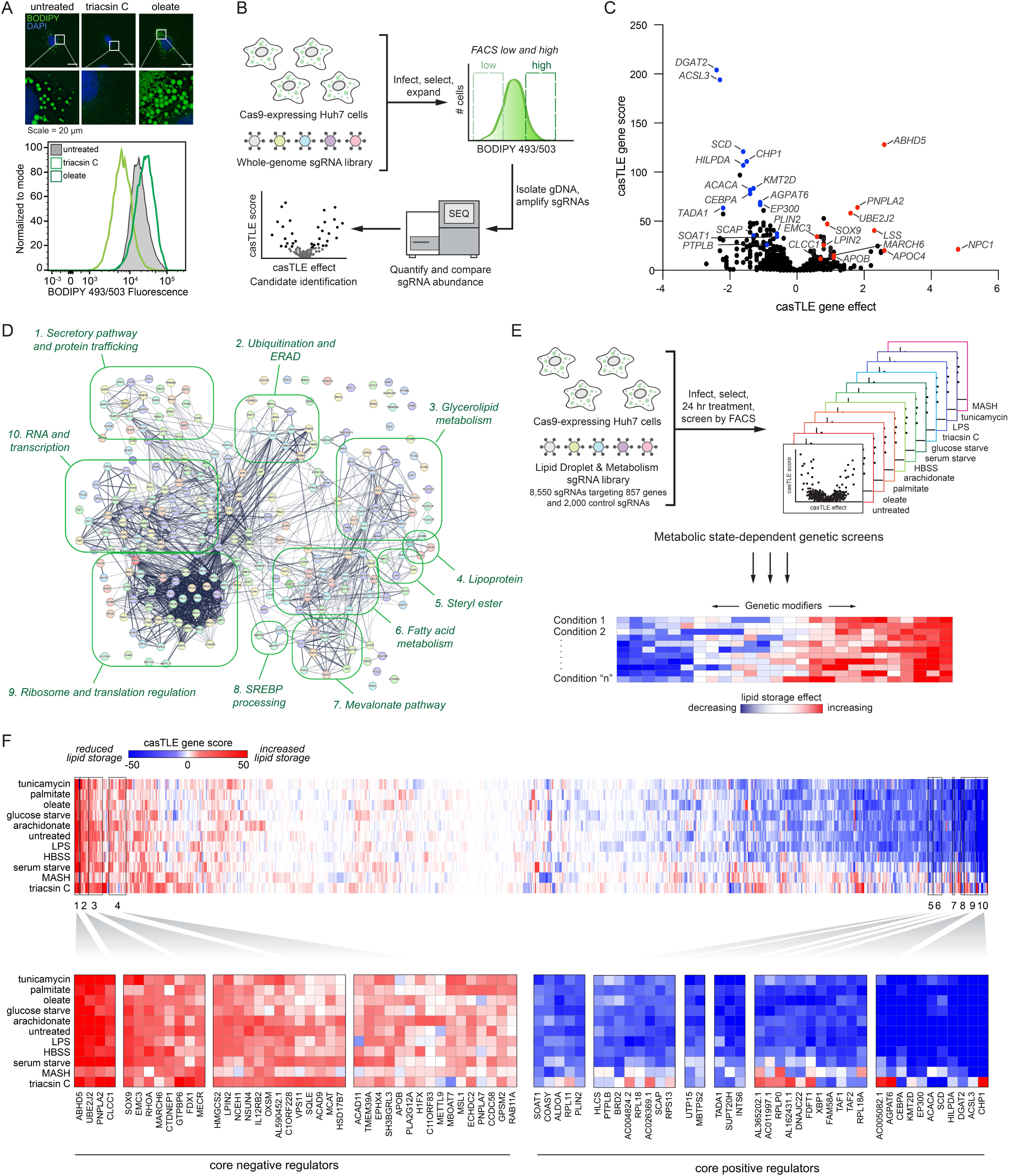
Parallel CRISPR-Cas9 screens identify metabolic state-dependent genetic modifiers of lipid storage. A) *Top*: Screen conditions were optimized in Huh7 cells treated for 24 h with 1 µg/mL triacsin C or 100 µM oleate and analyzed by fluorescence microscopy of cells labeled with BODIPY 493/503 (LDs) and DAPI (nuclei). *Bottom*: Flow cytometry BODIPY 493/503 fluorescence histograms of untreated, 1 µg/mL triacsin C- and 100 µM oleate-treated Huh7 cells. B) Schematic of FACS-based CRISPR-Cas9 screen approach to identify genes that regulate neutral lipid abundance, using BODIPY 493/503 as a neutral lipid reporter. C) Volcano plot indicating the gene effects (i.e., phenotype) and gene scores (i.e., confidence) for individual genes from batch retest screens in Huh7 cells. Gene effects and scores are calculated from two biological replicates. Positive (red) and negative (blue) genes of interest are highlighted. D) 265 of the 285 credible hits mapped in STRING confidence (text mining, experiments, physical interactions, genetic interactions, functional pathways) grouped manually by GO functional annotations. E) Schematic of parallel CRISPR screens under eleven different metabolic stress conditions. F) Heatmap of clustered genes based on gene score across all conditions. Boxes 1-4 indicate clusters of core positive regulators and boxes 5-10 indicate clusters of core negative regulators.

Most of our understanding of LD regulation is based upon studies in unperturbed and oleate-treated cells. However, the functions of LDs are influenced by diverse conditions and certain regulators are metabolic state specific^9^. To provide insights into the genetic modifiers of neutral lipid storage under different metabolic states, we performed duplicate FACS-based screens using our VLDM sgRNA library under 11 different metabolic conditions, including nutrient starvation, nutrient excess, and cell stress conditions (**Fig 1E and Supplementary Table 2**). Importantly, this series of 22 genetic screens generated extensive chemical-genetic interaction data that can be used to cluster genes with similar functional profiles in an unbiased manner, thereby facilitating functional predictions for novel candidate regulators (**Fig 1F**). The findings highlight the key importance of the glycerolipid metabolic pathways and identify differences in the utilization of members of enzyme families, such as the use of different *AGPAT*, *LIPN*, and *DGAT* enzymes under different conditions (**Extended Data Fig 2A and Supplementary Table 2**). Unbiased clustering revealed metabolic state-specific as well as core positive and negative regulators that generally reduced LDs and increased LDs, respectively, when the gene was depleted (**Fig 1F and Extended Data Fig 2B-K**). Genes that are known to play similar roles were often observed to cluster together, such as *BSCL2* (also known as *seipin*) and its binding partner *TMEM159* (also known as *LDAF1* and *promethin*) (**Extended Data Fig 2D**) which participate in LD nucleation and emergence from the ER^19^. The clustered core positive regulators included genes with roles in promoting neutral lipid synthesis (e.g., *ACSL3*, *DGAT2*, *CHP1*, *AGPAT6*, *SCD*), sterol ester synthesis (e.g., *SOAT1*), SREBP signaling (e.g., *SCAP*, *MBTPS2*), and LD stabilization (e.g., *PLIN2*) (**Fig 1F**). Conversely, the clustered core negative regulators included genes with roles in neutral lipid breakdown (e.g., *ABHD5*, *PNPLA2*), cholesterol ester breakdown (e.g., *NCEH1*), ER-associated degradation (ERAD) (e.g., *MARCH6*, *UBE2J2*), and lipoprotein secretion (e.g., *apoB*) (**Fig 1F**). These data establish a phenotypic-rich compendium of genetic modifiers of neutral lipid storage under multiple metabolic conditions. To promote accessibility and analysis, the data from these screens will be deposited in CRISPRlipid^9^, an online community driven data commons for functional genomics data related to lipid biology.

### Enlarged lipid droplets accumulate in cells lacking CLCC1

Many candidate regulators with no known roles in lipid metabolism and LD biology were identified (**Fig 1F and Supplementary Table 2**). CLIC-like chloride channel 1 (CLCC1) emerged as a priority candidate for characterization because: *1*) it clustered strongly with well-known lipolysis regulators *ABHD5* and *PNPLA2* (**Fig 1F and Fig 2A**), *2*) genetic variants in *CLCC1* in humans are associated with alterations in serum lipids (**Extended Data Fig 3**), and *3*) it was one of the most enriched genes among the core negative regulators with strong enrichment of multiple sgRNAs targeting *CLCC1* across several conditions (**Fig 1F and Extended Data Fig 4A**). *CLCC1* Huh7 knockout (CLCC1^KO^) cells generated with two independent sgRNAs increased lipid storage under multiple conditions (**Extended Data Fig 4B**). LDs in the CLCC1^KO^ cells were larger in size and they were stable following treatment with the acyl-CoA synthetase inhibitor triacsin C, which triggers the lipolytic consumption of LDs in control cells (**Fig 2B-D**). We also observed larger LDs in the CLCC1^KO^ cells by electron microscopy (EM) (**Fig 2E**). The CLCC1^KO^ cells had a ∼1.5-2-fold increase in TAG by thin layer chromatography (TLC), but no significant change in cholesterol esters (CE) **(Fig 2F)**. Employing a pulse chase assay that uses a fluorescently labeled fatty acid^9,20^, we found that loss of CLCC1 increased TAG biosynthesis and decreased TAG breakdown (**Extended Data Fig 4C,D**), suggesting that alterations of multiple aspects of neutral lipid metabolism contribute to the buildup of enlarged LDs in CLCC1^KO^ cells.

**Figure 2.**
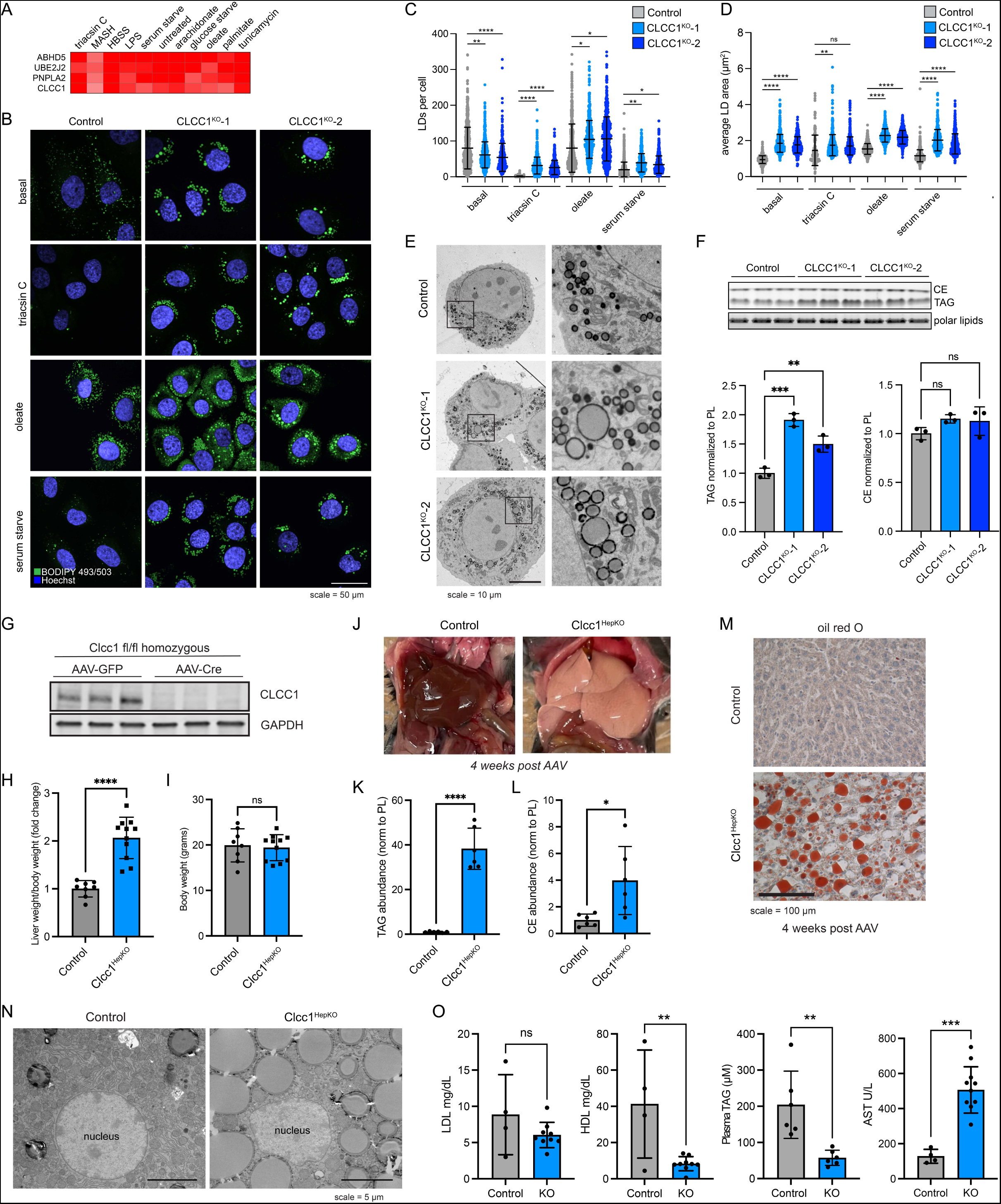
Loss of CLCC1 results in lipid droplet accumulation and hepatic steatosis. A) Cluster of top negative regulators of lipid storage from metabolic state-dependent CRISPR- Cas9 screens. B) Representative confocal images of lipid droplets using BODIPY 493/503 in control (expressing safe targeting sgRNA) and CLCC1^KO^ cells under basal conditions or following treatment with 1 µg/mL triacsin C for 24 h, 100 µM oleate for 24 h, or serum starve for 48 h. C) Quantification of the number of LDs from (C). Data represent mean ± SD of > 100 cells across three biological replicates. *\*\*\*\*p<0.0001 by one-way ANOVA with Dunnett’s multiple comparisons test*. D) Quantification of the area of LDs from (C). Data represent mean ± SD of > 100 cells across three biological replicates. *\*\*\*\*p<0.0001 by one-way ANOVA with Dunnett’s multiple comparisons test*. E) Representative transmission EM images of negative stained Huh7 cells expressing a safe targeting sgRNA (control) or sgRNAs against CLCC1. F) TLC resolving of TAG, CE, and polar lipids in Huh7 control and CLCC1^KO^ cells. Quantification of TAG (left graph) and CE (right graph) bands normalized to phospholipids. Data represent mean ± SD of three biological replicates. *\*\*\*\*p<0.0001 by one-way ANOVA with Dunnett’s multiple comparisons test*. G) Immunoblot analysis of three Clcc1 fl/fl mice injected with either AAV-GFP (control) or AAV- Cre (Clcc1^HepKO^). Samples were analyzed four weeks post-injection. H) Fold change in liver weight normalized to body weight for control and Clcc1^HepKO^ mice. *n > 9*. I) Body weight of the indicated control and Clcc1^HepKO^ mice. *n > 9*. J) Representative images of livers of control and CLCC1^HepKO^ m K,L) Quantification of TAG (K) and CE (L) normalized to PL using TLC. Data represent mean ± SD of six mice. *\*\*\*\*p<0.0001 by one-way ANOVA with Dunnett’s multiple comparisons test*. M) Representative oil red O stained liver sections from control and Clcc1^HepKO^ mice. N) Representative transmission EM images of negative stained control and Clcc1^HepKO^ m. O) Quantification of AST, LDL and HDL from clinical analyzer. Data represent mean ± SD of > four mice. *\*\*\*\*p<0.0001 by one-way ANOVA with Dunnett’s multiple comparisons test*.

### Deletion of *Clcc1* in mouse liver causes steatosis

*Clcc1* deletion is embryonic lethal in mice^21^. A spontaneous recessive mutation in *Clcc1* that disrupts the mouse *Clcc1* gene causes ER stress and cerebellar degeneration^22^ and the conditional knockout of mouse *Clcc1* in motor neurons also leads to ER stress and neurodegeneration^23^. However, the role of *Clcc1* in most tissues is unknown. To determine the physiological importance of liver *Clcc1*, floxed *Clcc1* mice were injected with either AAV8-Cre to knockout *Clcc1* in hepatocytes (Clcc1^HepKO^) or with AAV8-GFP as a control. Four weeks after the injection, we confirmed depletion of Clcc1 protein by immunoblotting (**Fig 2G**). Clcc1^HepKO^ mice had a ∼2-fold increase in liver weight/body weight ratio compared to control mice (**Fig 2H**), but no change in overall body weight (**Fig 2I**). Gross analysis of liver pathology revealed enlarged, whitened livers in the Clcc1^HepKO^ mice **(Fig 2J)**, indicative of lipid accumulation and hepatic steatosis. Indeed, TLC indicated a dramatic increase in TAG (**Fig 2K**) and CE (**Fig 2L**), and neutral lipid accumulation in the Clcc1^HepKO^ livers was evident based on oil red O staining (**Fig 2M**). EM provided additional evidence indicating the accumulation of enlarged LDs in the Clcc1^HepKO^ mice (**Fig 2N**). Finally, analysis of plasma indicated a reduction in TAG and HDL in the Clcc1^HepKO^ mice (**Fig 2O**), indicating a defect in hepatic lipid secretion. Despite the loss of Clcc1 for only four weeks and the lack of a nutritional challenge (e.g., high fat diet), an increase in AST, a marker of liver damage, was observed in the Clcc1^HepKO^ mice (**Fig 2O**). ALT was also measured, but there was no significant change (**Extended Data Fig 4E**). These findings demonstrate that Clcc1 plays an important role in regulating hepatic lipid storage and protecting hepatocytes from lipotoxicity.

### Lipid droplets in hepatocytes lacking CLCC1 are trapped within the ER lumen

As a first step towards understanding the mechanistic basis for LD accumulation in CLCC1^KO^ cells, we performed a pairwise comparison of our previously published PLIN2-GFP^9^ and current BODIPY 493/503 batch retest screens using the focused LD and metabolism-targeted sgRNA libraries (**Fig 3A**). As anticipated, there was a strong positive correlation in genetic modifiers identified in these two screens. However, CLCC1 was an outlier, with CLCC1 sgRNAs associated with an increase in neutral lipids, but a counter intuitive decrease in PLIN2-GFP (**Fig 3A**). This was surprising because the perilipin family of LD “coat” proteins are constitutively present on LDs^24^ and the levels of PLIN2 generally correlate with LD abundance^9,25^. Moreover, PLIN2-GFP is typically considered to be an obligate LD protein, and its degradation by the proteasome is induced under low LD conditions^9,26–28^. Western blotting confirmed that endogenous PLIN2 protein levels were undetectable in CLCC1^KO^ cells (**Fig 3B**). Incubation with the proteasome inhibitor MG132 rescued PLIN2 levels in the CLCC1^KO^ cells, indicating that despite high amounts of LDs, PLIN2 is being degraded post-translationally by the proteasome (**Extended Data Fig 5A**). Immunofluorescence staining of PLIN2 also revealed a strong reduction in PLIN2-positive LDs in the CLCC1^KO^ cells, with the large LDs appearing to be completely devoid of any PLIN2 staining (**Fig 3C**). Importantly, overexpression of CLCC1 rescued the PLIN2-GFP localization to LDs (**Extended Data Fig 5B**) and the reduction in PLIN2-GFP levels (**Extended Data Fig 5C**) in CLCC1^KO^ cells, consistent with the altered PLIN2 and LDs reflecting on-target depletion of *CLCC1*. Furthermore, proteomics analyses of LD-enriched buoyant fractions validated the reduction in PLIN2 on LDs and revealed the reduction in numerous well known LD proteins, such as ATGL, ABHD5, FSP1, LDAH, PNPLA3, and others (**Fig 3D and Supplementary Table 3**). These results indicate that although CLCC1^KO^ cells appear to accumulate large LDs, the LDs lack canonical LD proteins. To examine the generalizability of this phenotype, we generated knock out CLCC1 HepG2 (hepatoma) and U-2 OS (osterosarcoma) cell lines that were previously genetically engineered to express PLIN2 fused to GFP (i.e., PLIN2-GFP) at is genomic locus^9^. As in the Huh7 CLCC1^KO^ cells, HepG2 CLCC1^KO^ cells exhibited enlarged PLIN2-negative LDs (**Extended Data Fig 5D**), a reduction in PLIN2-GFP levels (**Extended Data Fig 5F**), and an increase in monodansylpentane (i.e., neutral lipid) staining (**Extended Data Fig 5F**). In contrast, LDs in the U-2 OS CLCC1^KO^ cells were of the expected size and were PLIN2-positive (**Extended Data Fig 5E**), and PLIN2 and neutral lipid amounts were also unchanged (**Extended Data Fig 5G**). These data suggest that the observed LD phenotypes are specific to hepatocytes.

**Figure 3.**
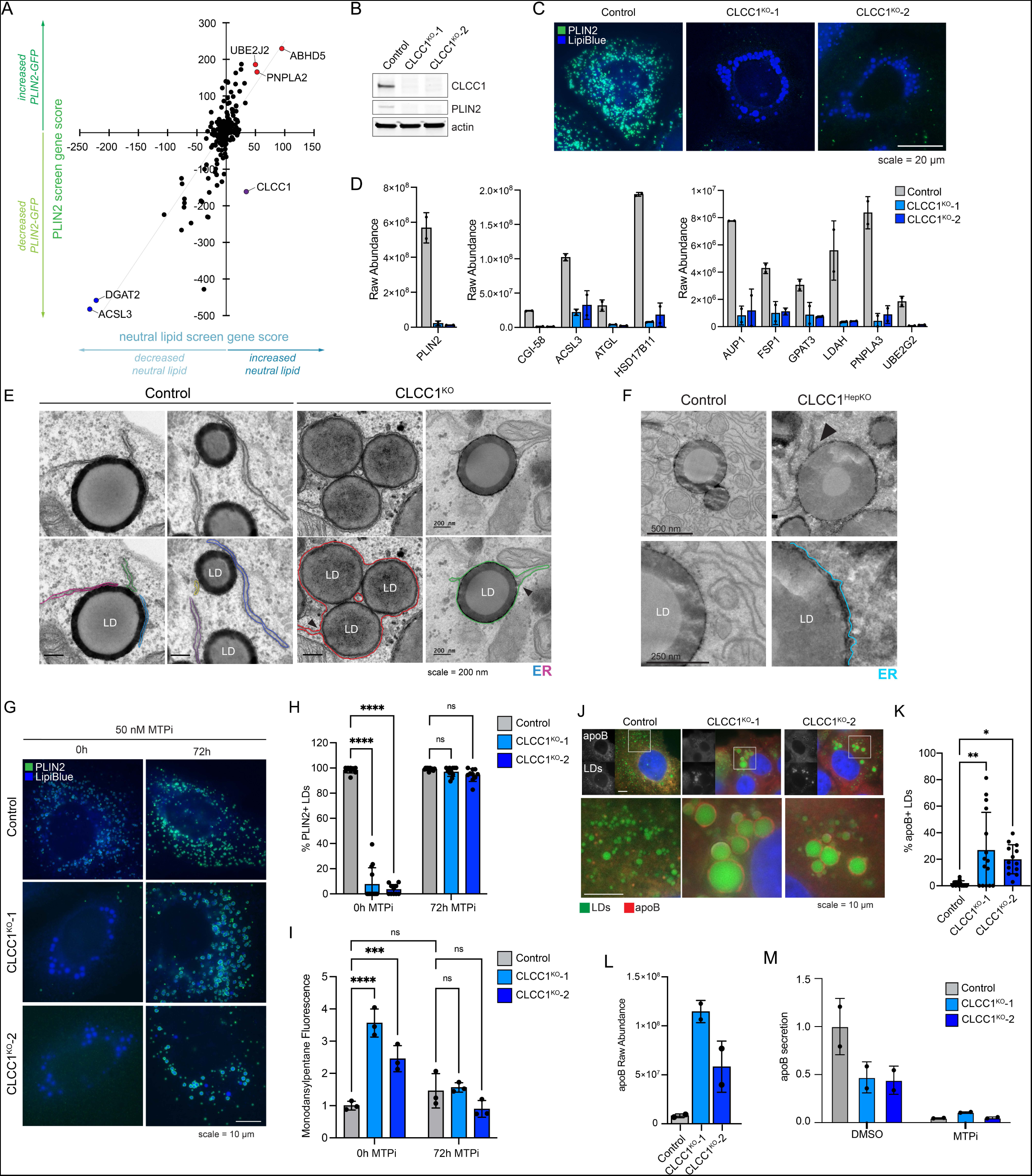
LDs in CLCC1^KO^ cells are trapped within the ER lumen. A) Pairwise comparison of the BODIPY 493/503 neutral lipid batch retest screen (Fig 1C and Supplementary Table 1) and a prior PLIN2-GFP batch retest screen^9^. B) Immunoblot analysis of PLIN2 levels in control and CLCC1^KO^ cells. C) Representative fluorescence microscopy images of PLIN2 and LDs in CLCC1^KO^ cells. PLIN2 was labeled with rabbit anti-PLIN2 antibody (green) and LDs (blue) were stained with 500 nM Lipi-Blue. Scale bar represents 20 µm. D) Raw abundance values for selected known LD proteins from buoyant fraction proteomics across two technical replicates. E) Transmission EM of negative stained control and CLCC1^KO^ cells. ER is highlighted in color and bifurcations, locations where the ER separates and a bilayer is found to encircle LDs, are denoted with black arrows. Scale bar represents 200 nm. F) Transmission EM of negative stained control and Clcc1^HepKO^ mouse liver. ER is highlighted in color and bifurcations are denoted with black arrows (as in panel E). Scale bar represents 200 nm. G) Representative fluorescence microscopy images of cells treated with the MTP inhibitor 50 nM CP-346086 for 72 h. Cells were fixed, PLIN2 was labeled with rabbit anti-PLIN2 antibody (green), and LDs were stained with 500 nM Lipi-Blue (blue). Scale bar represents 10 µm. H) Quantification of PLIN2-positive LDs in (C) where n > 10 cells. I) Quantification of flow cytometry measuring neutral lipid storage by monodansylpentane (i.e., Autodot) fluorescence. J) Representative fluorescence microscopy images of control and CLCC1^KO^ cells. ApoB was labeled with goat anti-apoB antibody (red) and LDs were stained with 500 nM Lipi-Blue (blue). Scale bar represents 10 µm. K) Quantification of the percentage of apoB-positive LDs in (H) where n > 10 cells. L) Raw abundance values for apoB from buoyant fraction proteomics across two technical replicates. M) Quantification of secreted apoB in medium using ELISA.

One potential mechanism that could explain the reduction of LD proteins is that the LD surface is somehow occluded, preventing protein trafficking to or insertion into the LD bounding monolayer. For example, tight organelle contacts can exclude proteins^29^. We therefore examined the distribution of several organelles and their relationship with LDs in the CLCC1^KO^ cells by immunofluorescence. The distribution and morphology of the Golgi, lysosomes, and mitochondria were mostly unchanged (**Extended Data Fig 6A**). However, the ER (marked by BFP-KDEL) exhibited an atypical morphology (**Extended Data Fig 6A,B**). Instead of sheets and tubules, strong ER staining was observed encircling the enlarged LDs in CLCC1^KO^ cells (**Extended Data Fig 6A,B)**. To further understand the relationship between the ER and LDs we performed EM. Typical ER-LD contact sites were observed between the ER and LDs in control cells (**Fig 3E**). In the CLCC1^KO^ cells, LDs were surrounded by a bilayer membrane (**Fig 3E**). The monolayer of the LD was distinguishable from the bounding bilayer (**Fig 3E)**. In some cases, we observed an ER sheet that was connected to a bilayer encircling one or more LDs (**Fig 3E**), indicating that LDs are present within the ER lumen instead of the cytoplasm. We observed a similar relationship in EM images of liver tissue from the Clcc1^HepKO^ mice, with LDs inside an encapsulating bilayer membrane (**Fig 3F**). These data indicate that LDs in CLCC1^KO^ cells are devoid of cytoplasmic LD proteins because they emerged into the ER lumen and are spatially segregated from cytoplasmic LD proteins. These findings further suggest that the LDs in CLCC1^KO^ cells are resistant to lipolysis because they are within the ER, rendering them inaccessible to ATGL and other cytoplasmic lipases.

### Lipid droplets in CLCC1^KO^ cells are MTP-dependent

The accumulation of PLIN2-negative LDs in the ER lumen was specific to hepatocyte-derived cell types (**Fig 3A-F, Extended Data Fig 5D-G)**, suggesting that a distinct property of hepatocyte lipid metabolism is important. A unique aspect of hepatocytes is their ability to form and secrete lipoproteins (i.e., VLDLs) to provide lipids to other tissues. Similar to LDs, lipoproteins consist of a neutral lipid core encircled by a phospholipid membrane decorated with regulatory proteins such as apolipoprotein B (apoB). The initial formation of lipoproteins involves microsomal triglyceride transfer protein (MTP)-mediated transfer of TAG into the ER lumen^11^. Incubation of CLCC1^KO^ cells with the MTP inhibitor CP-346086 rescued the biogenesis of PLIN2-positive LDs (**Fig 3G,H**) and the amount of neutral lipids (marked by monodansylpentane) (**Fig 3I**). MTP inhibitor treatment also rescued PLIN2 protein levels (**Extended Data Fig 6C**). There was no change in MTP levels in CLCC1^KO^ cells (**Extended Data Fig 6D**). In addition, we observed an increased percentage of apoB positive LDs in the CLCC1^KO^ cells (**Fig 3J,K**), though not all LDs were apoB positive. There was also an increased amount of apoB peptide counts in buoyant, LD-enriched fractions (**Fig 3L**). Consistent with the reduced HDL and TAG measured in the mouse serum of Clcc1^HepKO^ mice (**Fig 2O**), we observed a reduction in apoB secretion into medium by CLCC1^KO^ cells (**Fig 3M)**. No change in the levels of secreted albumin in medium was observed (**Extended Data Fig 7A**), suggesting that secretion is not generally disrupted. In addition, a luciferase-based assay of ER protein folding and secretion, which uses the substrate asialoglycoprotein receptor 1 (ASGR1) fused to luciferase Cluc^30,31^, also showed no impairment in the CLCC1^KO^ cells (**Extended Data Fig 7B**). Together, these findings indicate that the enlarged LDs are aberrant lipoproteins that exhibit reduced secretion. The near complete lack of cytoplasmic PLIN2-positive LDs suggests a profound shift in neutral lipid flux away from cytoplasmic LDs towards lumenal MTP-dependent lipoproteins. It is noteworthy that VLDLs typically have a diameter of ∼50-80nm, whereas the mean diameter of the lumenal LDs in the CLCC1^KO^ cells is ∼1.84 µm. Thus, the lipoproteins that accumulate in the CLCC1^KO^ cells are exceptionally large in volume compared to a normally secreted VLDL particle (>10,000-fold larger).

### CLCC1 and its relationship with ER stress and ER scramblases

Loss of CLCC1 has been associated with an increase in ER stress^22,23^. However, there was no increase in the mRNA transcripts of a variety of common ER stress targets (**Extended Data Fig 7C**) or the protein levels of the ER chaperone grp78/BiP (**Extended Data Fig 7D**) in CLCC1^KO^ cells. Moreover, although induction of ER stress by treatment with tunicamycin and thapsigargin altered LD distribution, the LDs remained PLIN2-GFP positive (**Extended Data Fig 7E**). These findings indicate that ER stress is not necessary for the lumenal LDs CLCC1^KO^ cells nor is it sufficient to trigger the accumulation of lumenal LDs.

*CLCC1* was the only gene that exhibited an increase in neutral lipid amount and a decrease in PLIN2-GFP in the pairwise comparison of our batch retest screens (**Fig 3A**). Broader examination of our genome-wide screens revealed that depletion of *TMEM41B* was also associated with an increase in neutral lipids and a decrease in PLIN2-GFP (**Extended Data Fig 8A**). TMEM41B is an ER scramblase that interacts with apoB and regulates hepatic lipoprotein secretion^32^. *TMEM41B* KO in mice reduces lipoprotein secretion and results in severe hepatic steatosis and the accumulation of LDs encapsulated by ER membranes^32^. The KO of a second ER scramblase, VMP1, causes hepatic steatosis and the buildup of PLIN2-negative LDs, similarly suggesting an association between phospholipid scrambling and ER lumenal LDs in hepatocytes^33,34^. However, in our screens in Huh7 cells, loss of *VMP1* was associated with a reduction in PLIN2-GFP but no change in neutral lipid staining (**Extended Data Fig 8A**).

There was no change in the levels of TMEM41B or VMP1 in CLCC1^KO^ cells (**Extended Data Fig 8B**). To explore the relationship of CLCC1 and ER scramblases, we generated *TMEM41B* and *VMP1* KO Huh7 cells (**Extended Data Fig 8C,D**). LDs were PLIN2 positive (i.e., cytoplasmic) in the VMP1^KO^ cells (**Extended Data Fig 8E**). A portion of LDs in TMEM41B^KO^ cells were enlarged and PLIN2 negative (**Extended Data Fig 8E**), consistent with an ER lumenal localization. However, in contrast to CLCC1^KO^ cells, TMEM41B^KO^ cells exhibited PLIN2 crescent staining (**Extended Data Fig 8E**), which suggests that a fraction of LDs in the TMEM41B^KO^ cells are trapped in the membrane with half of the LD facing the cytoplasm and the other half the ER lumen. In addition, although MTP inhibition completely rescued LD cytoplasmic emergence in the CLCC1^KO^ cells (**Fig 3G,H**), MTP inhibition only had a partial effect on TMEM41B^KO^ cells and the effect was not significant (**Extended Data Fig 8F,G**). Finally, overexpression of either TMEM41B or VMP1 in CLCC1^KO^ cells had no effect on the amount of PLIN2-negative and -positive LDs (**Extended Data Fig 9A-C**). These data indicate that while there are similarities in the phenotypes and a potential functional relationship, dysregulation of ER scramblases alone was not sufficient to account for the altered neutral lipid flux in the CLCC1^KO^ cells.

### CLCC1 is structurally homologous to yeast Brl1p / Brr6p

CLCC1 was first identified through a series of blast searches using a partial sequence of a yeast chloride channel Mid-1^35^. It was noted that CLCC1 exhibits no overall sequence similarity with Mid-1 and no similarity with known channel families^35^. There was, however, a short motif in a CLCC1 transmembrane domain that was partially present in the CLIC family of chloride channels, leading to the naming of CLCC1 initially as Mid-1-related chloride channel (MCLC) and eventually as CLCC1^35^. Altered chloride conductance in ER microsomes isolated from cells overexpressing CLCC1 has been observed^23,35^, but direct evidence for ion conductance using purified CLCC1 is lacking. Moreover, the predicted alpha-fold structure of CLCC1 is considerably different from canonical CLIC chloride channels (**Extended Data Fig 10A,B**), suggesting that if CLCC1 conducts chloride it would be through a novel mechanism. Given these data, we considered the possibility that CLCC1 may have alternative biochemical functions.

An HHpred search for remote homologs revealed a strong relationship between amino acids 204-378 of CLCC1 with the yeast paralogs Brl1p (probability score 94%) and Brr6p (probability score 86%) (**Fig 4A and Extended Data Fig 11**). This is based on both sequence and structural homology. Indeed, the alpha-fold predicted structures showed a homology domain with very similar features (**Fig 4B**). Both predicted structures contain an elongated transmembrane helix (TMH) within a portion of a helix that enters deep into the lumen, a sharp turn followed by a short helix (knuckle region), a perpendicular amphipathic helix (AH), and another TMH. These features are absolutely conserved in both families. The knuckle region of Brl1p and Brr6p contains two cysteine pairs that stabilize the structure with intramolecular disulfide bonds^15^, and CLCC1 contains one cysteine pair. Co-immunoprecipitation data with differentially tagged CLCC1 indicated that CLCC1 self-associates^23^, possibly forming a dimer or higher order oligomer. Indeed, proteome-scale predictions of homo-oligomeric assemblies predict a CLCC1 dimer that is stabilized by intermolecular disulfides in a lumenal N-terminal domain not shared with Brl1p / Brr6p^36^ (**Fig 4C, Extended Data Fig 12A**). Consistent with this model, we find that in the absence of reducing agent, CLCC1 migrates as a dimer (**Fig 4D**). The addition of DTT yields the expected CLCC1 monomer (**Fig 4D**). Furthermore, analysis of the CLCC1 Brl1p homology domain using ColabFold suggests the possibility of larger CLCC1 oligomers that form a ring structure (**Extended Data Fig 12B, Supplementary Videos 1-3**). The same is also the case for Brl1p and Brr6p (**Extended Data Fig 12C, Supplementary Videos 4-6,** and data not shown).

**Figure 4.**
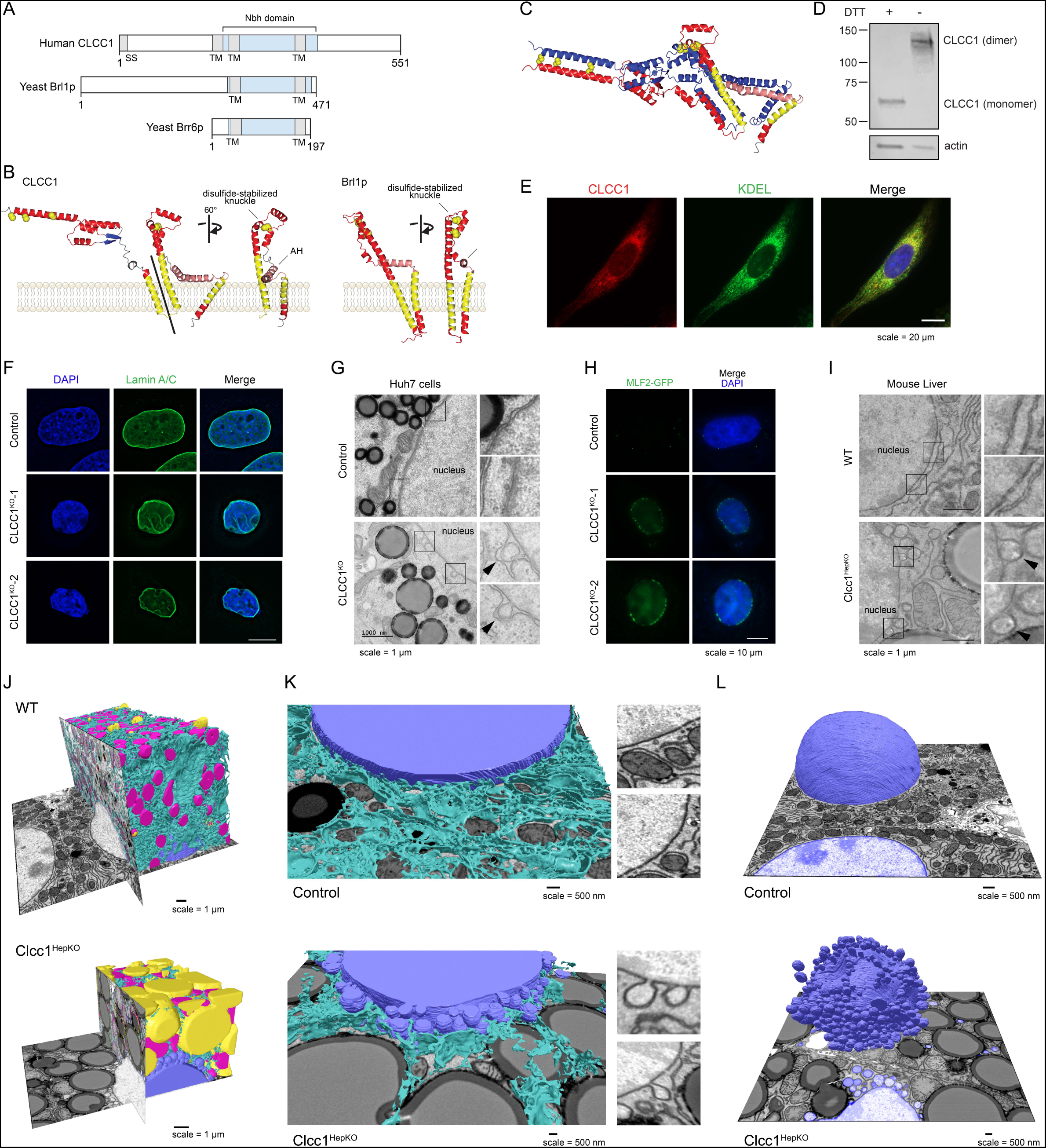
CLCC1 mediates membrane remodeling and is the human homolog of yeast Brl1p. A) Domain structure of human CLCC1, yeast Brl1p, and yeast Brr6 indicating transmembrane domains (TM), signal sequence (SS), and the homologous ‘NPC-biogenesis “h”-shaped (Nbh) domain’ in blue. B) Human CLCC1 and yeast Brl1p homologous region alpha-fold structure. Ribbon colors: yellow =TMH, pink = AH, red = other helix; blue = sheet; yellow spheres = conserved cysteines. C) Predicted disulfide-stabilized CLCC1 dimer. Colors: one protomer as in B, one all blue. D) Western blot analysis of CLCC1 in the presence and absence of the reducing agent DTT. E) Fluorescence microscopy images of CLCC1 (red), the ER marker KDEL (green) and nuclei (DAPI, blue) in control and CLCC1^KO^ cells. F) Representative fluorescence microscopy images of nuclei (DAPI, blue) and lamin A/C (green) in control and CLCC1^KO^ cells. G) Transmission EM of negative stained control and CLCC1^KO^ cells. Zoomed regions are included to highlight the NE ultrastructure. H) Representative fluorescence microscopy images of the NE herniation marker MFL2-GFP (green) and DAPI-stained nuclei (blue). Scale bar represents 20 µm. I) Transmission EM of negative stained control and Clcc1^HepKO^ mouse liver. Zoomed regions are included to highlight the NE ultrastructure J,K) Partial (J,K) reconstruction of segmented raw FIB-SEM data with ER (cyan), mitochondria (magenta), nuclei (blue) and LDs (yellow) from hepatocytes of WT (above) and Clcc1^HepKO^ (below) mice. Insets in panel K show examples of nuclear membrane in the WT and Clcc1^HepKO^ mice. L) Partial reconstruction of segmented raw FIB-SEM data with nuclei (blue), from hepatocytes of WT (above) and Clcc1^HepKO^ (below) mice.

### CLCC1 plays a critical role in nuclear pore assembly

Brl1p and Brr6p function in the assembly of nuclear pore complexes, which are large nuclear envelope (NE) protein complexes that perforate the inner and outer NEs to enable regulated exchange between the nucleus and cytoplasm^13–18^. Although the detailed architecture of nuclear pore complexes has been determined^37^, their mechanisms of biogenesis and insertion into membranes remains incompletely understood. *BRL1* and *BRR6* are essential, and conditional disruptions result in nuclear membrane herniations (also referred to as nuclear blebs) indicative of disruptions in nuclear pore complex insertion, as well as altered membrane composition and synthetic interactions with lipid metabolic pathways^13–18^. It is noteworthy that there is precedence for factors that have roles in both nuclear pore complex assembly and hepatic neutral lipid storage. The ER lumenal AAA ATPase torsinA and its cofactors Lull1 and Lap1 promote nuclear pore complex biogenesis, and loss of these factors results in nuclear membrane herniations^38^. In addition, the conditional loss of torsinA, Lull1, or Lap1 in mouse liver causes hepatic steatosis and reduced lipoprotein secretion^39,40^.

Given the structural homology between CLCC1 with yeast Brl1p / Brr6p, we examined a potential relationship of CLCC1 with nuclear structure, including nuclear morphology and the integrity of the NE in CLCC1^KO^ cells. Interestingly, co-essentiality analyses^41^ indicate a functional relationship between CLCC1 and nuclear pore complex genes (**Extended Data Fig 13A**). In addition, proteomic analyses of isolated organelles^42^ indicate that CLCC1 is present in the ER, with similar localization profiles as the torsin activators Lap1 (also known as TOR1AIP1) and Lull1 (also known as TOR1AIP2) as well as the scramblases TMEM41B and VMP1 (**Extended Data Fig 13B**). Consistent with these data and past studies^21,22^, fluorescence imaging revealed that endogenous CLCC1 localized to both the ER and NE (**Fig 4E**). Fluorescence imaging of CLCC1^KO^ cells indicated alterations in lamin A/C staining and nuclear morphology (**Fig 4F, Extended Data Fig 14**). Importantly, EM showed nuclear membrane herniations in the cultured CLCC1^KO^ cells (**Fig 4G**), similar to those found following depletion of Brl1p and Brl6p in yeast^13–18^. NE herniations occur when there is a failure of inner and outer NE fusion during nuclear pore complex insertion, resulting in membrane protrusions that contain extruded nucleoplasm. Consistent with this structure, puncta of GFP-tagged myeloid leukemia factor 2 (MLF2), a marker of NE herniations that accumulates in the nucleoplasmic interior of the NE herniation^43^, decorated the NE of CLCC1^KO^ cells but not control cells (**Fig 4H**). Similar to our cultured cells, extensive NE herniations were also observed in the liver tissue of CLCC1^HepKO^ (**Fig 4I**). KO of *CLCC1* in the osteosarcoma cell line U-2 OS led to the accumulation of NE herniations (**Extended Data Fig 14B**), despite the absence of lumenal LDs in this cell line. Furthermore, although MTP inhibition rescues LD biogenesis (**Fig 3G-I**), MTP inhibition had no effect on the NE herniations in the CLCC1^KO^ cells (**Extended Data Fig 14C,D**). We also observed NE herniations in the TMEM41B^KO^ cells, though they were less abundant than in the CLCC1^KO^ cells (**Extended Data Fig 14E,F**). Together, these data indicate that while the effect of *CLCC1* KO on neutral lipid flux is restricted to hepatocytes, which secrete lipoproteins, there is a more generalizable role for CLCC1 in nuclear pore complex assembly across multiple cell lines. These data also indicate that the NE herniations are not downstream of the defects in neutral lipid flux.

To characterize the impact of Clcc1 on organelle architecture, we performed focused ion beam scanning EM (FIB-SEM) analyses of liver tissue from control and Clcc1^HepKO^ mice (**Fig 4J-L, Extended Data Fig 15A,B, and Supplementary Videos 7-8**). As anticipated, this analysis revealed the accumulation of numerous enlarged LDs that occupied a large percentage of the cell volume, at the expense of other organelles (**Fig 4J and Extended Data Fig 15C**). One of the most remarkable phenotypes in the Clcc1^HepKO^ liver tissue was the presence of extensive NE herniations decorating the majority of the nucleus (**Fig 4K,L and Extended Data Fig 15D**). Most of the nuclear NE herniations are connected to the NE, often by a thin membrane stalk (**Fig 4K,L**). A small subset of blebs was not observed to have a NE connection, perhaps indicating shedding of the bleb into the cytoplasm (**Fig 4L**). We also observed large indentations in the nucleus that were generated by juxtanuclear LDs pressing into the nucleus and that had reduced of amounts NE herniations (**Fig 4L and Extended Data Fig 15D**). Together, our findings implicate CLCC1 as the Brl1p / Brr6p human homolog that promotes nuclear pore complex assembly.

## Discussion

In the current study, we performed a series of CRISPR-Cas9 screens under diverse metabolic conditions, establishing a phenotypically rich compendium of hepatic neutral lipid storage genetic modifiers. The chemical-genetic interactions present in our compendium of lipid storage regulators identified CLCC1 as an important mediator of lipid storage. Indeed, we find that loss of CLCC1 results in the accumulation of LDs and severe hepatic steatosis in mice. The lumenal location of LDs, requirement for MTP, and immunostaining with apoB suggest that these LDs are aberrant lipoproteins. It is noteworthy that these lumenal LDs accumulate at the expense of cytoplasmic LDs, indicating that loss of CLCC1 causes a profound shift in ER neutral lipid flux.

We identify a role for CLCC1 in nuclear pore complex assembly (**Fig 5**). In metazoan cells with open mitosis, the NE disassembles to enable chromosome segregation and then reassembles around chromatin^18^. Cells that employ open mitosis form nuclear pore complexes through two mechanisms, either by forming nuclear pore complexes on membranes associating with chromatin prior to the reformation of a sealed NE or through inside-out insertion of the nuclear pore complex into the intact NE during interphase^18^. The inside-out insertion mechanism is similar to the sole mechanism of nuclear pore complex biogenesis employed by organisms with closed mitosis, such as yeast^18^. This process involves the initial insertion of a set of nuclear pore complex subunits into the inner NE, causing the NE to dimple and thereby reducing the distance between the inner and outer NE membranes^18^. Through a poorly understood process, the inner and outer nuclear membranes fuse, perforating the NE and enabling the completion of a mature nuclear pore complex^18^. The discovery of Brl1p, Brr6p, and their binding partner Apq12p as NE assembly factors in yeast was an important breakthrough^13–18^. However, traditional BLAST searches failed to identify a metazoan homolog of these proteins, leading to the proposal that this mechanism was restricted to organisms with closed mitosis. How nuclear membrane fusion during nuclear pore complex insertion occurs in metazoan cells has remained a mystery. Our data solve this mystery and implicate CLCC1 as the long sought Brl1p / Brr6p human homolog. This conclusion is supported by sequence conservation and structural homology in terms of multiple features, as well as evidence of shared functions in nuclear pore complex assembly. Indeed, similar to loss of Brl1p and Brr6p, *CLCC1* KO results in nuclear membrane herniations, reflecting a failure in inner and outer nuclear envelope fusion during nuclear pore complex biogenesis.

**Figure 5.**
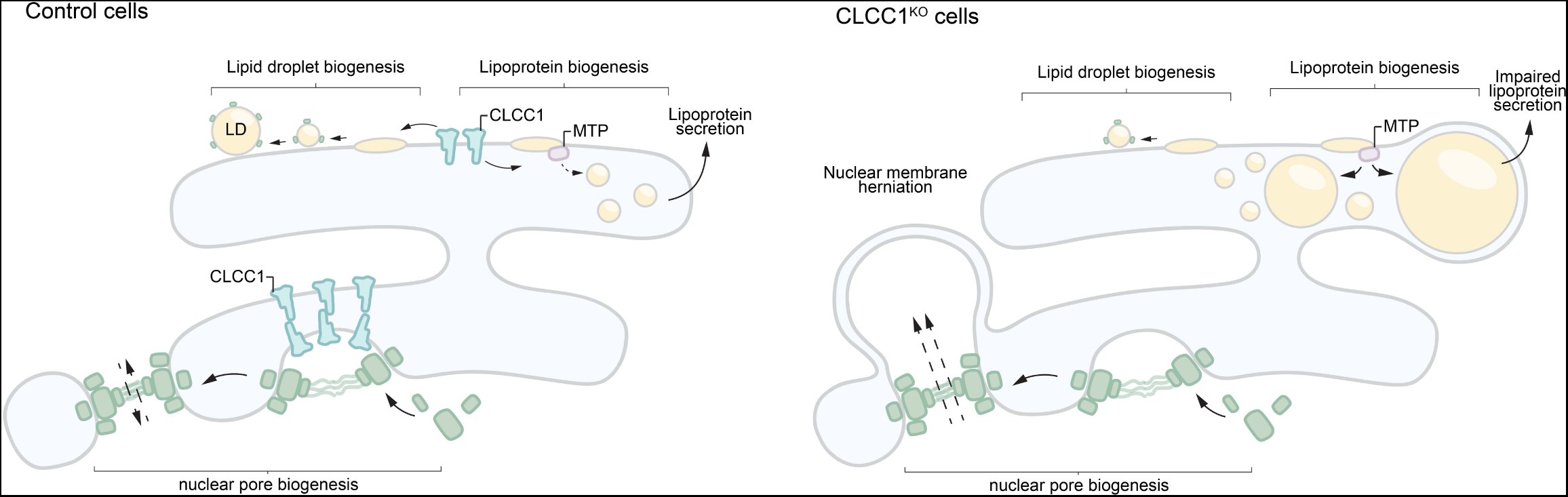
CLCC1 mediates membrane remodeling and is the human homolog of yeast Brl1p. Model depicting the cellular functions of CLCC1 and the impact of CLCC1 depletion. CLCC1 is an ER and NE multi-pass transmembrane protein that promotes nuclear pore complex assembly and is required for the fusion of the inner and outer NE membranes. In the absence of CLCC1, nuclear pore complex assembly is impaired and large nuclear membrane herniations (i.e., blebs) accumulate. CLCC1 also regulates neutral lipid flux between cytoplasmic LDs and lumenal lipoproteins. CLCC1^KO^ cells exhibit reduced neutral lipid channeling into cytoplasmic LDs and instead neutral lipid are channeled into enlarged MTP-dependent lipoproteins that build up in the ER lumen and fail to be properly secreted. (Also see **Extended Data Fig 16**)

The integral membrane portion of CLCC1 and Brl1p / Brr6p (TMH1-Knuckle-AH-TMH2) is a conserved domain. We propose the name ‘NPC-biogenesis “h”-shaped (Nbh) domain’, based on the shared functions and shape which is reminiscent of a lowercase “h” (**Fig 4A**). One conserved feature of Nbh domains is the separation of TMH1 and TMH2 by the lumenal AH, which leads to the TMHs being unconstrained by each other, so that they need not be parallel. This allows Nbh proteins to reside in curved membranes. A second feature is that all of TMH1, AH and TMH2 are strongly predicted by Alphafold Multimer [and similar algorithms - same results from DMFold and Multifold] to contain extensive interfaces that oligomerize side-by-side. Finally, the whole domain has a triangular cross-section, as shown by predicted dimers with the AHs pointing in different directions (**Fig 4B**) and overall shape (**Extended Data Fig 12A**), which is the basis upon which ring oligomers can be predicted^36^. These features together provide the first explanation at a molecular/physical level for how Brl1p / Brr6p impact nuclear pore complex biogenesis, which has been missing since they were first linked genetically to this process^14,44^. Our protein structure models suggest a potential mechanism in which a CLCC1 homo-oligomer locally bends the membrane due to the TMHs positioned at increasingly extreme angles to the perpendicular, which also applies to the yeast homologs (**Extended Data Fig 12B,C**). We were able to obtain ring structures between 12-20 protomers, with the major difference being the size of the central pore (1.4-5.8nm). Determining the stoichiometry of this putative oligomer is an important research direction that could shed light on the CLCC1 mechanism of action. Prior modeling suggests the possibility of trans interactions between Brl1p and/or Brr6p on opposing inner and outer NE membranes, which is supported by a prediction of a Brl1p-Brr6p heterodimer that interact via their knuckles (**Extended Data Fig 12D**)^45^. The putative CLCC1 oligomer would be predicted to bend the membrane inward, and interactions with a second CLCC1 homo-oligomer on an opposing membrane could facilitate membrane fusion (**Extended Data Fig 16**).

Our data implicate CLCC1 in the control of neutral lipid flux and lipoprotein biogenesis (**Fig 5**). One possibility is that CLCC1 could provide a lipid salvage pathway using a similar membrane fusogenic function as in NPC assembly, but with hemifusion of the ER membrane to a lumenal LD’s single leaflet. This would reconnect lumenal LDs with the inner leaflet of the ER and allow neutral lipids to regain the ability to be stored in cytosolic LDs through Ostwald ripening and emergence from the cytosolic leaflet (**Extended Data Fig 16, bottom**), a process which may require ER scramblases. Our findings, along with the emerging data examining the roles of torsinA complexes reveals a striking connection between a set of factors involved in nuclear pore complex assembly and hepatic neutral lipid flux. The precise role of torsinA in nuclear pore complex assembly and neutral lipid flux is unknown, but it is tempting to speculate that torsinA, CLCC1, and TMEM41B could function together, perhaps with CLCC1 oligomers as substrates of torsinA^39,40^. This is similar in concept to other AAA ATPases being dedicated to a very narrow range of targets, such as NSF regulation of the recycling of membrane fusogens (i.e., SNARE proteins). The similar phenotypes observed in cells lacking ER scramblases also suggests a potential functional role, such as in regulating NE membrane curvature or providing phospholipids for cytosolic LD emergence. Although many questions remain for future studies, our findings make an important advance by identifying a role for CLCC1 in hepatic neutral lipid flux and nuclear pore complex assembly.

## AUTHOR INFORMATION

Correspondence and requests for materials should be addressed to J.A.O. and A.P.A.

## ACKNOWLEDGEMENTS

This research was supported by grants from the National Institutes of Health to J.A.O. (R01GM112948 and R01DK128099), a grant from Merck Sharp & Dohme LLC, a subsidiary of Merck & Co., Inc., Rahway, NJ, USA to J.A.O., and a predoctoral fellowship from the National Science Foundation to A.J.M. J.A.O. is a Chan Zuckerberg Biohub Investigator. S.P.P. was supported by PREP Scholar grant from NIH (R25GM140276). T.P.L was supported by the Biotechnology and Biological Sciences Research Council, UK (BB/P003818/1). Thank you to the staff at the University of California Berkeley Electron Microscope Laboratory for advice and assistance in electron microscopy sample preparation and data collection. We thank S. Eyles (UMass Amherst, RRID: SCR_019063) for assistance with high-resolution mass spectrometry acquired on an Orbitrap Fusion mass spectrometer funded by NIH Grant 1S10OD010645-01A1. FIB-SEM data collected using the Zeiss XB 550 was supported NIH S10 Grant S10OD030258.

## AUTHOR CONTRIBUTIONS

A.J.M. and J.A.O. conceived of the project, designed the experiments, and wrote the majority of the manuscript. All authors read, edited, and contributed to the manuscript. A.J.M. and K.K.D. performed the proteomics and A.J.M. and M.L. the thin layer chromatography. A.J.M, C.B, E.T., E.S.M., K.K.D., and M.A.R. contributed to cell line generation and analysis. A.J.M. and M.L. performed lipid measurements. G.K. and Y.Z. contributed to the generation of the floxed CLCC1 mice. E.S.M., E.T., S.P.P., A.P.A, and G.P. generated the knockout CLCC1 mice and contributed to the tissue and serum analyses. E.T., S.T., and Y.Z. assisted with serum analyses. A.J.M., D.M.J., R.Z., M.K., A.P.A, and G.P. performed tissue processing and electron microscopy. A.P.A, and G.P. performed segmentation and analyses of FIB-SEM data. S.T. and Y.Z. contributed intellectually to the development of the project throughout. T.P.L. performed the protein structure homology searches and structure modeling.

## COMPETING INTERESTS

J.A.O. is a member of the scientific advisory board for Vicinitas Therapeutics.

**Note added in proof:** During the preparation of this manuscript an additional, complementary characterization of CLCC1 in hepatic lipid flux was performed. Please refer to the corresponding preprint - Wu et al., “ CLCC1 Governs Bilayer Equilibration at the Endoplasmic Reticulum to Maintain Cellular and Systemic Lipid Homeostasis.”

## Methods

### Cell lines and culture conditions

Huh7, U-2 OS, and HEK293T cells were cultured in DMEM containing 4.5 g/l glucose and L-glutamine (Corning) supplemented with 10% FBS (Thermo Fisher Scientific and Gemini Bio Products), penicillin, and streptomycin. HepG2 cells were cultured in RPMI 1640 medium (Gibco) containing L-glutamine supplemented with 10% FBS, penicillin, and streptomycin. All cells were maintained at 37°C and 5% CO_2_.

Generation of endogenously labeled PLIN2-GFP Huh7, HepG2, and U-2 OS reporter cells are detailed in Roberts et al., 2023.

### Plasmids and cloning

All knockout cell lines (in Huh7, HepG2, and U-2 OS cells) were generated using the pMCB320 plasmid, a gift from M. Bassik (Addgene, 89359). Guide sequences for CLCC1, TMEM41B, VMP1, and safe-targets (sgSAFE #5784) (see Table 1) were selected from the Bassik Human CRISPR Knockout Library (Addgene, 101926, 101927, 101928, 101929, 101930, 101931, 101932, 101933, 101934). Guide sequences were cloned into pMCB320 using the restriction enzymes BstXI and BlpI. For exogenous protein expression, CLCC1 (DNASU, HsCD00951632), 1X FLAG (DYKDDDDK)-tagged CLCC1, TMEM41B (DNASU, HsCD00829148), and VMP1 (DNASU, HsCD00080545) were cloned using the Gibson assembly and the Gateway system (Thermo Fisher, 11791020) in a pLenti-CMV-Hygro vector (Addgene, 17454).

### Generation of CRISPR-Cas9 genome edited cell lines

To generate lentiviral particles, lentiCas-Blast plasmid (Addgene, 52962) was co-transfected with third-generation lentiviral packaging plasmids (pMDLg/pRRE, pRSV-Rev, and pMD2.G) into HEK293T cells. Lentiviral media was collected 72 hr after transfection, passed through a 40 µm filter, and then used to infect Huh7 (wild type, PLIN2-GFP), HepG2 (wild type, PLIN2-GFP), and U-2 OS (wild type, PLIN2-GFP) cells. Cells were selected in media containing blasticidin (4 μg/ml in Huh7 and U-2 OS; 6 μg/ml in HepG2) for 5 days. Active Cas9 expression was validated by flow cytometry analysis following infection with a self-cleaving <u>mCherry</u> plasmid (pMCB320 containing mCherry and an sgRNA targeting the mCherry gene).

Lentiviral particles with sgRNA-containing pMCB320 plasmids were generated as described above and used to infect cells stably expressing Cas9. After 72 hr of growth, infected cells were selected in media containing puromycin (2 μg/ml in Huh7 and HepG2; 1 μg/ml in U-2 OS) until over 90% cells were mCherry positive and all uninfected control cells were dead. Huh7 CLCC1^KO^ and TMEM41B^KO^ clones (in wild type and PLIN2-GFP backgrounds) were isolated using serial dilutions. Knockout efficiencies were confirmed via immunoblotting.

### Genome-wide Huh7 CRISPR-Cas9 screens

All CRISPR-Cas9 screens reported here were described previously in Roberts et al., 2023 and in Mathiowetz et al., 2023. Genome-wide CRISPR-Cas9 screens were performed using the Bassik Human CRISPR Knockout Library. The library consists of nine sublibraries, comprising a total of 225,171 elements, including 212,821 sgRNAs targeting 20,549 genes (∼10 sgRNAs per gene) and 12,350 negative-control sgRNAs. Lentiviral particles containing each sublibrary were generated as described above. Huh7 cells stably expressing Cas9 were transduced with lentiviral packaged sublibraries (one sublibrary at a time) in 8 μg/ml polybrene. After 72 hr of growth, infected cells were selected in media containing 2 μg/ml puromycin until over 90% of cells were mCherry positive (via flow cytometry). Cells were then recovered for 3-5 days in media lacking puromycin and frozen in liquid nitrogen.

For the screen, library infected cells were thawed (one sublibrary at a time) and maintained at 1000x coverage (1,000 cells per library element) in 500 cm^2^ plates (about 20^6 cells per plate). Each library was passaged once before sorting. On the day of the sort, cells were dissociated using 0.25% Trypsin-EDTA (Gibco), collected by centrifugation at 300 x g for 3 min, stained with 1 µg/µL BODIPY 493/503 (Thermo Fisher, D3922) in DPBS on ice for 30 minutes, then washed 1x with DPBS. Cells were resuspended in phenol red free media (HyClone, 16777-406) supplemented with 3% FBS and 1% fatty acid-free BSA and kept on ice until FACS.

Cells were sorted on a BD FACS Aria Fusion equipped with 4 Lasers (488 nm, 405 nm, 561 nm, and 640 nm). sgRNA-expressing, mCherry+ cells were gated into the brightest 30% and dimmest 30% by the 488 nm laser. Cells were sorted into 15 ml conicals containing DMEM with 4.5 g/L glucose and L-glutamine supplemented with 10% FBS. For each sort, 1000X cells were collected (500X in each gate). Sorted cells were collected and sequenced according to Mathiowetz et al., 2023. Results from the genome-wide screen are available in Table 2.

### Lipid Droplet and Metabolism Library CRISPR-Cas9 screens

The custom human VLDM library contains 10,550 elements, with 8,550 sgRNAs targeting 857 genes (∼10 sgRNAs per gene) and 2,000 negative control sgRNAs. Guide sequences were from the Bassik Human CRISPR Knockout Library, and the library construction protocol and cell line generation were previously described.

For each screen, cells were thawed and expanded at >1000x coverage. For all screens, cells were seeded into 500 cm^2^ plates at 1,000-fold library coverage. For the Huh7 metabolic state-dependent screens, cells were treated the following day with 1) no treatment, 2) 1 μg/ml triacsin C for 24 hr, 3) 100 μM oleate-BSA complex for 24 hr, 4) HBSS (Gibco, 14025092) for 24 hr, 5) 0.2% FBS-containing DMEM (serum starve) for 48 hr, 6) glucose-free DMEM (Gibco, 11966025) for 24 hr, 7) 50 μM palmitic acid for 24 hr, 8) 5 μM arachidonic acid for 24 hr, 9) 5 μg/mL tunicamycin for 24 hr, 10) 500 ng/mL LPS for 24 hr, or 11) NASH stress mix for 16 hr. NASH stress mix was defined as 10 mM glucose, 5 mM fructose, 400 µM oleic acid, 200 µM palmitic acid, 100 ng/mL LPS, and 30 ng/mL TNFɑ. Cells were screened by FACS as described above. Results from each metabolic state screen are available in Table 3.

### CRISPR screen data analysis

Sequence reads were aligned to the sgRNA reference library using Bowtie 2 software. For each gene, a gene effect and score (likely maximum effect size and score) and *p*-values were calculated using the Cas9 high-Throughput maximum Likelihood Estimator (casTLE) statistical framework as previously described.

Morpheus (https://software.broadinstitute.org/morpheus/) was used to perform unbiased gene clustering on metabolic state screens. Genes were ranked according to casTLE score and complete Euclidean linkages. Functional interactions and protein-protein interactions for high confidence candidate regulators were identified using the STRING database^45^.

### General animal care

All experimental procedures involving animals were approved by the UC Berkeley Institutional Animal Care and Use Committee. Mice were maintained from 6 to 12 weeks of age on a 12 hr light:12 hr dark cycle at room temperature in the UC Berkeley pathogen-free barrier facility with free access to water and standard laboratory chow diet (PicoLab Mouse Diet 20 no. 5058, LabDiet). We used CLCC1 flox/flox and Albumin-Cre in the C57BL/6J genetic background (stock no. 000632). These animals were a generous gift from Dr. G. Ku at UCSF. Experimentation was performed between 8-12 weeks of age. In animal experiments, all measurements were included in the analysis. Mice were randomly allocated to groups; the only criteria were sex and age as explained above. The sample size and number of replicates for this study were chosen based on previous experiments performed in our laboratory and others. No specific power analysis was used to estimate sample size. Imaging studies could not be done blinded owing to the evident intrinsic features of the datasets. *In vivo* studies could not be blinded owing to the adenoviral injection protocol. Experimental and control samples were processed together using the same conditions.

### Floxed CLCC1 mouse generation

Clcc1 flox mice (generated by the Knockout Mouse Project) were obtained from the University of California Davis Mouse Biology Program (C57BL/6N-Atm1Brd Clcc1tm1a(KOMP)Mbp/JMmucd), where exon 7 has been floxed. The neomycin selection cassette and lacZ reporter were removed by breeding to CAG-Flpo (C57BL/6N-Tg(CAG-Flpo)1Afst/Mmcd). Mice were then bred for at least 4 generations to C57BL/6J animals, removing the CAG-Flpo. Genotyping for the flox-ed allele from genomic DNA was performed with the following PCR primers: TCATGACATGAACCATATGTGAATTCC and CACCATGCCTGGCTACAAATGC.

### Adenovirus-mediated deletion of CLCC1

To deplete CLCC1, 8-week old homozygous CLCC1-flox mice were injected with either AAV-Cre or AAV-GFP. 1.5 x 10^11^ GC of virus was diluted in 100 uL of PBS and injected via the tail vein. Mice were euthanized by CO_2_ 4 weeks post-injection. Livers were photographed. >4 mice were analyzed per experiment. The liver was weighed and divided into sections which were flash frozen in liquid nitrogen, transferred on dry ice, and stored a −80 °C.

### Mouse plasma collection and analysis

Blood was collected via submandibular vein puncture and centrifuged at 2,000 x g in Microtainer SST (BD, 365967) tubes for 15 minutes at 4 °C to isolate plasma. Plasma was analyzed via Clinical Analyzer. Triglycerides were quantified by a luciferase-based assay (Promega, J3160)

### Flow cytometry

Cells were washed 2x in DPBS, dissociated using TrypLE Express (Gibco, 12605010), collected by centrifugation at 500 x g for 5 min, and stained with 1 µg/µL BODIPY 493/503 or 200 µM monodansylpentane (MDH) (Abcepta, SM1000b) in DPBS on ice for 30 minutes.

For all flow cytometry assays, fluorescence was analyzed using an LSR Fortessa (BD Biosciences). The following filter sets were used: FITC (GFP, BODIPY 493/503), Pacific Blue (BFP, MDH), and Texas-Red (mCherry). FlowJo Software (BD Biosciences) was used to quantify fluorescence and generate representative histograms.

### Immunoblotting

Cells were lysed in 1% SDS and sonicated at 15% amplitude for 15 seconds. For albumin secretion, cells were incubated for 24 h in FBS-free DMEM, and proteins were precipitated from the media with acetone. Animal tissues were homogenized in 1% SDS with an immersion homogenizer for 15 seconds. Protein concentrations were determined and normalized using a BCA <u>protein assay</u> (Thermo Fisher Scientific, 23225). Equal amounts of protein by weight were combined with Laemmli buffer, boiled for 10 min at 95 °C, separated on 4–20% polyacrylamide gradient gels (Bio-Rad Laboratories) and transferred onto nitrocellulose membranes (Bio-Rad Laboratories). Membranes were incubated in 5% nonfat milk in PBS with 0.1% Tween-20 (PBST) for 30 min to reduce nonspecific antibody binding. Membranes were then incubated overnight at 4 °C in PBST containing antibodies diluted in 5% BSA, followed by incubation for at least 1 hr in fluorescence-conjugated secondary antibodies diluted in PBST containing 5% nonfat milk. Immunoblots were visualized on a LI-COR imager (LI-COR Biosciences), and Fiji/ImageJ was used for quantification of protein levels.

### Fluorescence microscopy

For widefield microscopy of live cells, Huh7, HepG2, and U-2 OS cells were grown in 4-well or 8-well Nunc™ Lab-Tek™ II Chambered Coverglass (Borosilicate Glass 1.5; Thermo Fisher Scientific, 155360) coated with poly-L-lysine. Lipid droplets were stained with 1 μM BODIPY 493/503 for 30 minutes or 500 nM LipiBlue for 30 min, nuclei were stained with 5 μg/mL Hoeschst 33342 (Thermo Fisher Scientific, 62249) for 30 min, lysosomes were stained with 75 nM Lysotracker DND-22 (Thermo Fisher Scientific, L7525) for 30 min, and mitochondria were stained with 500 nM Mitotracker Green (Thermo Fisher Scientific, M7514) for 30 min. For imaging the ER, cells were transiently transfect with BFP-KDEL (Addgene, 49150) and imaged 48 hr later. For imaging of nuclear blebs, cells were transiently transfected with MLF2-GFP (a gift from Dr. C. Schlieker) and imaged 48 hr later. Prior to imaging, cells were washed 2x with DPBS and imaged in fresh phenol red-free medium supplemented with 10% FBS. Live cells were imaged using a Zeiss Axio Observer 7 fitted with a 63X oil objective using DAPI, GFP, Cy-3, and Cy-5 filters. Cells were imaged at 37 °C with 5% CO_2_. Z-stacks of 0.2-μm thickness were acquired.

For widefield microscopy of fixed cells, Huh7 cells were grown in 12-well plates on glass coverslips coated with poly-L-lysine. Cells were washed 3x with DPBS, fixed for 15 min in 4% (w/v) PFA in DPBS and washed 3x again with DPBS. Cells were permeabilized for 15 min with 1% BSA in DPBS containing 0.1% Triton X-100 when staining for ER, Golgi, or nuclear proteins or 0.01% digitonin when staining for LD proteins and then washed 3x with DPBS. Cells were incubated in antibodies to PLIN2 (Abcepta, AP5118c), GM130 (Cell Signaling, 12480), CLCC1 (Thermo, HPA009087), KDEL (Abcam, ab276333), or lamin A/C (Cell Signaling, 4777) diluted at 1:1000 in 1% BSA in DPBS for 1 hr in the dark. Lipid droplets were stained with 1 μM BODIPY 493/503 for 30 minutes or 500 nM Lipi-Blue for 30 minutes, nuclei were stained with 1 µg/mL DAPI, and primary antibodies were blotted with fluorescent secondaries (Thermo Fisher, A21202, A-21109) diluted at 1:1000 for 30 min in the dark. Cells were washed 3x with DPBS and coverslips were mounted on 1 mm glass slides using Fluoromount-G (SouthernBiotech, 0100-01).

For live cell <u>confocal microscopy</u>, Huh7 cells were grown in 24-well glass bottom plates (170 μm coverglass bottom; Eppendorf, 0030741021; Cellvis, P24-1.5H-N). Cells were either untreated or incubated in 100 μM oleate-BSA complex for 24 hr, 1 μg/ml triacsin C for 8 hours, or 0.2% FBS-containing DMEM (serum starve) for 48 h. Lipid droplets were stained with 1 μM BODIPY 493/503 and nuclei were stained with 5 μg/mL Hoeschst 33342 for 30 minutes. Prior to imaging, cells were washed 2x with DPBS and imaged in fresh phenol red-free medium supplemented with 10% FBS. Live cells were imaged using an Opera Phenix Plus High-Content Screening System (Perkin Elmer) confocal microscope equipped with a 40X water immersion objective using DAPI and GFP filters. Cells were imaged at 37 °C with 5% CO2. Z-stacks of 0.3-μm slices were acquired.

Images were merged and brightness and contrast adjusted using Fiji/ImageJ (https://imagej.net/software/fiji/). LDs were quantified by creating a custom analysis sequence using Harmony High Content Image Analysis Software, v4.9 (Perkin Elmer). For each field, maximum projection Z-stacks were processed with advanced flatfield correction. Nuclei and cytoplasm were defined using the DAPI and GFP channels, respectively, and border cells were automatically excluded from analyses. LDs were defined using the “Find Spots” building block (Lipi-Green stain, GFP channel), thresholding for size, intensity, and roundness. For each cell, lipid droplet number and area were quantified. LD quantification data were graphed and analyzed in Prism 9 (GraphPad).

### Transmission electron microscopy

For cell lines, Huh7 and U-2 OS cells were grown on 3 cm gridded LabTek dishes and fixed in 2% paraformaldehyde and 2% glutaraldehyde in PBS. Samples were stained with 1% osmium tetroxide + 1.5% potassium ferrocyanide for 1 hr and 1% uranyl acetate overnight. The next day, samples were washed and subsequently dehydrated in grades of ethanol (10 min each; 30%, 50%, 70%, 95% and 2 × 10 min at 100%). Samples were embedded in increasing concentrations of eponate resin mixed with ethanol (30 min each; 1:2, 1:1, 2:1 and 100% acetone) followed by polymerization in 100% eponate overnight at 50 °C.

For liver tissues, mice were anesthetized with 300 mg/kg ketamine and 30 mg/kg xylazine in PBS and perfused with 10mL of DPBS followed by 10mL of fixative buffer containing 4 parts of FP stock (2.5 % PFA, 0.06 % picric acid in 0.2M Sodium Cacodylate buffer pH 7.4) and 1 part of 25 % glutaraldehyde. After perfusion, small pieces (1–2 mm^3^) of liver were sliced at 300-micron thickness with a compresstome, transferred into a fresh fixative solution containing and incubated at 4C overnight. Samples were then washed in ice-cold 0.15 M sodium cacodylate buffer for 5 min, 3 times, and then incubated in 0.1 M sodium cacodylate solution containing 1% osmium tetroxide and 1.5% potassium ferrocyanide for 1 hr at 4 °C. Samples were rinsed 3x with water and incubated for 20 min in 1% thiocarbohydrazide and rinsed again 3x for 5 min with water. Samples were incubated in 2% OsO4 for 30 min and then rinsed 3x for 5 min with water, followed by washing 3x and incubation overnight at 4 °C in 1% uranyl acetate in MB. The next day, samples were washed and subsequently dehydrated in grades of acetone (10 min each; 50%, 70%, 90% and 2 × 10 min at 100%). Samples were embedded in increasing concentrations of eponate resin mixed with acetone (30 min each; 50%, 70%, 90% and 100% acetone) followed by incubation in 100% eponate for 4 hr. The samples were moved to fresh 100% eponate and polymerized at 65 °C for 24 hr.

The resin embedded sample blocks were trimmed, and 70 nm ultrathin sections were cut using a Leica UC6 ultramicrotome (Leica Microsystems, Vienna, Austria) and collected onto formvar-coated slot grids. Sections were imaged to find target regions using a Tecnai 12 120kV TEM (FEI, Hillsboro, OR, USA) and data recorded using an Gatan Rio16 CMOS camera and GMS3 software (Gatan Inc., Pleasanton, CA, USA).

### Structure predictions

Monomeric and multimeric sequences were submitted to AlphaFold2 using MMseqs2 using either the Google Colabatory^46^ or COSMIC2^47^ or were submitted to DMFold, MultiFOLD, or trRosetta^48^. The pLDDT of core homology regions as monomers and ring oligomers predicted by AlphaFold2: CLCC1 residues 209-353: monomer 79.2%, 16-mer: best 80.9%, average 80.2%; Brr6 residues 44-185: monomer 80.5%, 16-mer: best 78.1%, average 77.0%^49,50^.

### Focused ion beam scanning electron microscopy (FIB-SEM)

Mouse livers were fixed and prepared described above. The trimmed sample blocks were glued with silver paint (Ted Pella Inc.) onto Al stubs, and sputter coated (Pd/Au) with a Tousimis sputter coater on top of a Bio-Rad E5400 controller. Focused Ion Beam Scanning Electron Microscopy (FIB-SEM) imaging were performed using a Zeiss Crossbeam 550 (Carl Zeiss Microsystems GmbH, Oberkochen, Germany). The sample was tilted at 54° in order to perpendicular to ion beam. The FIB milling and SEM imaging of the target area were set up using Atlas 5 3D tomography (Carl Zeiss Microsystems GmbH, Oberkochen, Germany). Slices with a thickness of 10 nm were milled from the target area using the 30 kV 300pA ion beam. Energy-selective Backscattered (ESB) images were collected at 1.5 kV 1nA, with a dwell time of 18 ns, image pixel size of 10 nm, and tilt correction angle of 36°. The collected images were aligned with the Slice Registration in Dragonfly 2022.2 (Comet Technologies Canada Inc., Canada).

### FIB-SEM data segmentation, quantification, and visualization

Ground truth labels were generated by manually annotating each class (ER, mitochondria, nucleus, and lipid droplets) in five consecutive full-size images using Napari (v0.4.18). Tunable 2D-U-Net networks (DLSIA) were used to obtain rough predictions for each class^51^. These rough predictions were manually proofread and corrected in Napari. A block consisting of at least 250x250x250 voxels was used to train and fine-tune 3D-U-Net network models with Incasem^52^. Additional proofreading and manual corrections were performed in Napari. Objects, images, videos and quantifications from each class were generated using Arivis Vision 4D (v3.6.0).

### Liver histology

Liver preparation was performed as described above with 4% PFS. Liver pieces were flash frozen, embedded in OCT, and cryosectioned into 10 µm-thick sections. Liver sections were fixed in 4% paraformaldehyde for 20 min and stained with either oil red O. Images were acquired on a Zeiss LSM 880 FCS.

### BODIPY 558/568 C12 incorporation assay

Huh7 safe-targeting control and CLCC1^KO^ cells were seeded in 60-mm plates at 350 cells per plate. To determine the rate of LD biogenesis, cells were incubated in BODIPY C12-BSA complex (complete media + 1% BSA + 1 μM BODIPY 558/568 C12) for 0, 1, 3, or 6 hr. Cells were harvested by washing them twice, collecting in cold DPBS, and transferring to Eppendorf tubes (Eppendorf, 022363352). Cells were centrifuged at 500 x g for 5 min, washed in DPBS, and centrifuged again. Cell pellets were stored at −80 °C until the lipid extraction step. For the lipolysis assay (measuring loss of esterified C12), cells were incubated in BODIPY C12-BSA complex for 16 hr. Cells were then washed 3x with media and incubated in fresh media for 1 hr. Cells were then treated with 1 µg/mL triacsin C for 0, 6, or 24 hr. Cells were harvested, and pellets stored at −80 °C as described above.

### Triglyceride measurements by thin layer chromatography

Cell pellets were thawed at room temperature and resuspended in 50 μL DPBS. Liver tissues (approximately 30 mg, three per mouse) were homogenized in 1 ml methanol using an immersion homogenizer for 5 min at 4 °C. Lipids were extracted by adding tert-butyl methyl ether (1250 μL) and methanol (375 μL). The mixture was incubated on an orbital mixer for 1 hr at room temperature. To induce phase separation, water (315 μL) was added, and the mixture was incubated on an orbital mixer for 10 min at room temperature. Samples were centrifuged at 1,000x g for 10 minutes at room temperature. The upper organic phase was collected and subsequently dried *in vacuo*.

Dried lipid extracts were reconstituted in 30 µL (cells) or 200 µL (liver) chloroform/methanol (2:1, v/v). Lipids were then separated using HPTLC Silica gel 60 F254 plates (Sigma, 1137270001). 10 µL of the cell samples and 2 µL of the liver samples were spotted onto TLC plates and developed in CHCl3/EtOH/TEA/H2O (5:5:5:1, v/v). Plates were imaged on a ChemiDoc MP Imaging System (Bio-Rad Laboratories). Band densitometry analysis was performed using Image Lab 5.0 (Bio-Rad Laboratories). The reported mean ± standard deviation was determined from three biological replicates.

### Proteomic Analysis of LD Proteins

Safe-targeting and CLCC1^KO^ cell lines were grown until confluent in 500 cm^2^ plates of cells were scraped harvested in DPBS, centrifuged for 10 min at 500 x g, and stored at −80 °C. Buoyant fractions containing 1% SDS were acidified to a final concentration of 15% TFA. Samples were then cooled on ice for 30 min and centrifuged at 20,000 x g for 30 min at 4°C. The protein pellet was washed 3 times with 500 μL of ice-cold acetone and centrifuged for 10 min between each wash. The protein pellet was then dried in a vacuum evaporator for 10 min. Dried, precipitated proteins were resuspended in 0.1% RapiGest with 6 μL of sequencing-grade trypsin (Promega, 0.5 μg/μL) added to each sample and digested overnight at 37°C. Trypsinized samples were quenched with a final concentration of 5% TFA. Samples were desalted using the Waters Sep-pak 1cc (50mg) C18 cartridge.

Peptides were resuspended in 1% formic acid and 0.5 μg of peptides were separated on an Easy nLC 100 UHPLC equipped with a 15 cm nano-liquid chromatography column. Using a flow rate of 300 nL/min, the linear gradient was 5% to 35% over B for 90 min, 35% to 95% over B for 5 min and 95% hold over B for 15 min (solvent A 0.1% formic acid in water, solvent B 0.1% formic acid in ACN). Peptide identified and relative abundances were determined using Proteome Discoverer 2.4. Results are represented as average ± s.d. of duplicates.

### ApoB ELISA assay

Safe-targeting and CLCC1^KO^ cells were seeded in 6 cm plates and treated with 1 µg/mL DMSO or 50 nM MTPi for 72 hr. 24 hr before harvesting, cells were changed into FBS- and phenol red-free media. Media was collected and ApoB ELISA Assay (Sigma Aldrich, RAB069) was performed according to protocol. ApoB levels were normalized to cell protein levels and results are represented as average ± s.d. of biological duplicates.

### ASGR Luciferase Assay

The ASGR reporter plasmid was generated by Dr. Hotamisligil’s lab, as previously described^30,31^. Safe-targeting and CLCC1^KO^ cells were infected with lentivirus containing the ASGR construct. For the experiment, cells were changed to a fresh medium containing phenol red-free and incubated for 24 h with or without increasing doses of thapsigargin. 10 µl of media was transferred to 96-well white plates (Corning) for luciferase assays following the manufacturer’s protocol. Briefly, 50 µl of luciferase substrate (1 µM Cypridina (CLUC) or 10 mM CTZ (GLUC) in 100 mM tris buffer, pH 7.5) was added to the 10 µl medium and incubated in the dark for 5 min. The luminescence was read on Infinite 200 PRO plate reader (Tecan).

### Statistical analysis with Prism

All statistical analyses were performed using Prism 9 (GraphPad). For each panel, the number of biological replicates (n), *p*-values, and statistical tests employed are reported in figure legends and methods.

### Materials availability

All unique/stable reagents generated in this study are available from the <u>lead contact</u> with a completed Materials Transfer Agreement. The custom Human VLDM sgRNA Library will be deposited at Addgene.

## Data availability

Data available upon request from the lead contact.

## Extended Data Figure Legends

**Extended Data Figure 1.**
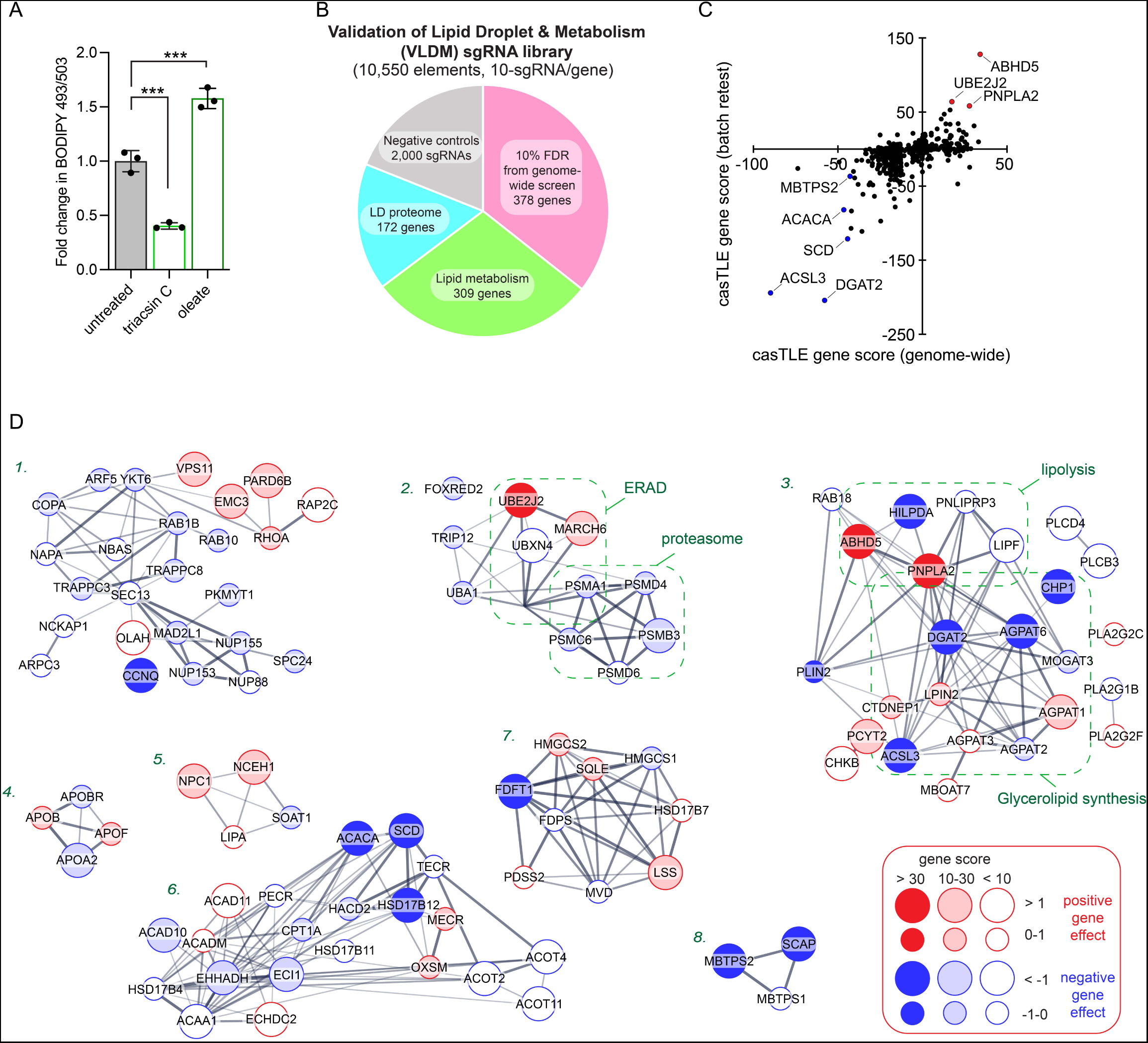
Analysis of lipid storage CRISPR-Cas9 screens. A) Quantification of the amount of neutral lipid from flow cytometry histograms (representative histograms shown in Fig 1A). Data represent mean ± SD across three biological replicates. *\*\*\*\*p<0.0001 by one-way ANOVA with Dunnett’s multiple comparisons test*. B) Breakdown of the custom lipid metabolism CRISPR-Cas9 library used in batch retest screens. C) Pairwise comparison of gene scores between the genome-wide and batch retest screens. Scores were adjusted as positive or negative based upon the direction of gene effect. D) Significant gene clusters from Figure 1D. Nodes are marked based on directionality of effect (red or blue), gene effect size, and confidence score.

**Extended Data Figure 2.**
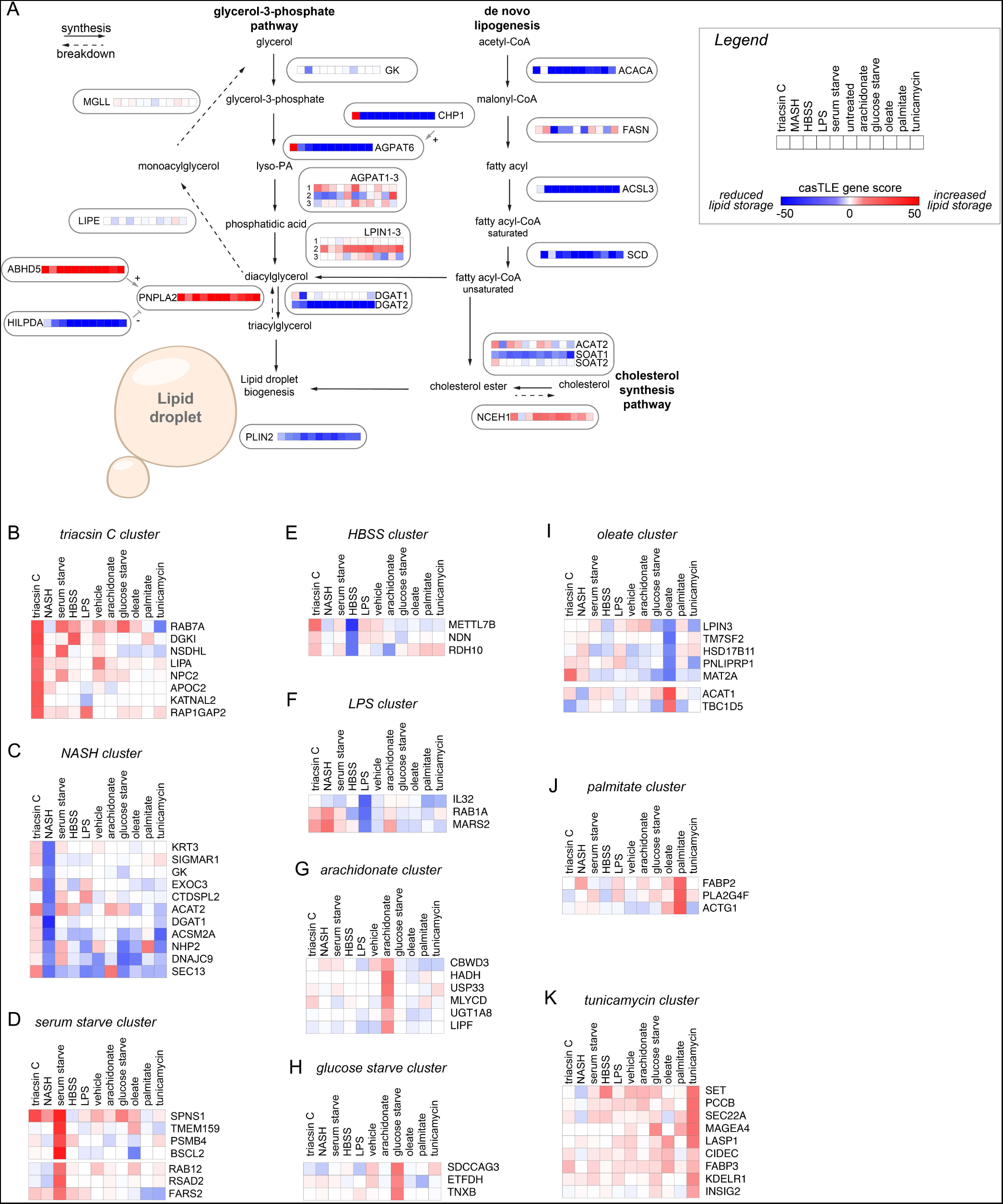
Enriched pathways and metabolic state-specific regulators of lipid storage. A) Schematic of LD synthesis and breakdown pathways. Genes are annotated with heatmaps corresponding to the gene score across metabolic conditions. Metabolic states follow the order indicated in the legend. B-K) Selected examples of gene clusters exhibiting metabolic state specific effects on neutral lipid storage.

**Extended Data Figure 3.**
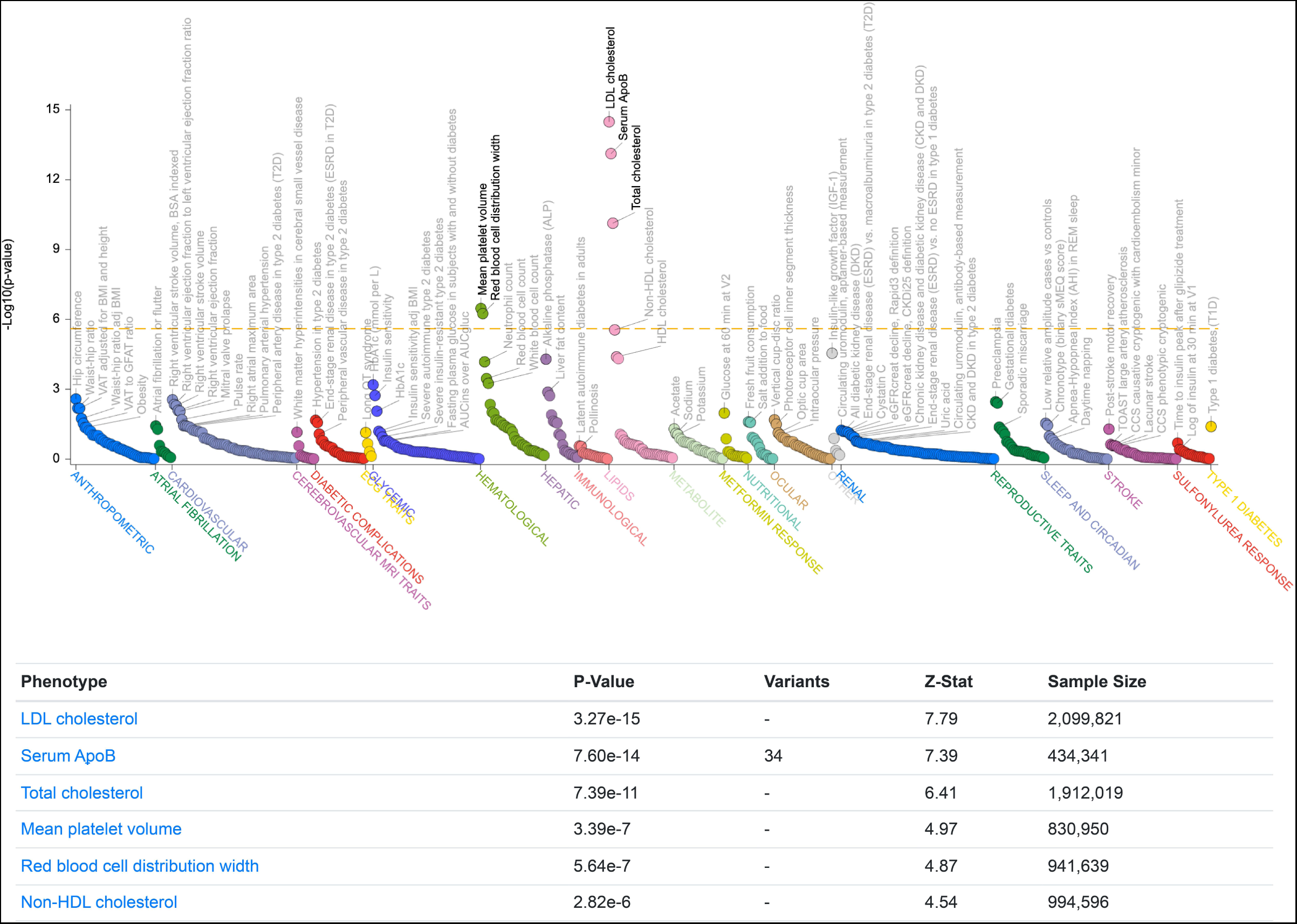
Genetic association of CLCC1 variants with altered serum lipids. Common variant gene-level associations for CLCC1 from the Common Metabolic Diseases Knowledge Portal. The plot shows phenotypic associations for CLCC1 based upon genetic associations using Multi-marker Analysis of GenoMic Annotation (MAGMA)^53^.

**Extended Data Figure 4.**
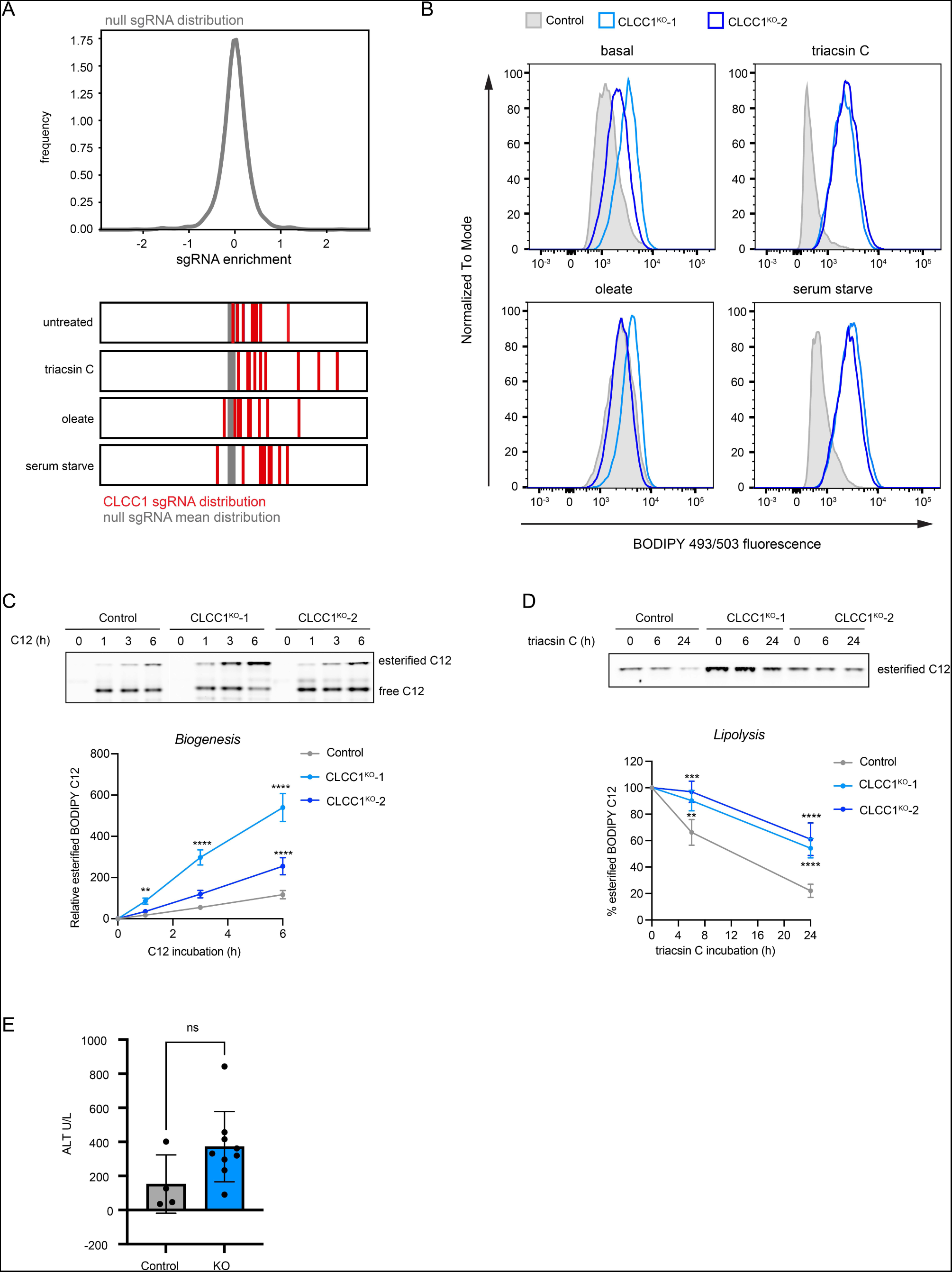
Loss of CLCC1 results in the accumulation of neutral lipids. A) Histogram indicating the distribution of control (null) sgRNAs. Shown below is the relative enrichment of CLCC1 targeting sgRNAs (red lines) under untreated, triacsin C, oleate, and serum starve conditions. The gray line indicates the mean of the control sgRNA distribution. B) Representative flow cytometry histograms of control and CLCC1^KO^ Huh7 cells following treatment with 1 µg/mL triacsin C for 24 h, 100 µM oleate for 24 h, serum starve for 48 h, or basal levels. C) Representative TLC resolving esterified and free BODIPY C12 558/568 in Huh7 cells expressing a safe targeting sgRNA or sgRNAs against CLCC1. Cells were incubated with BODIPY C12 for the indicated times, followed by lipid extraction and TLC. A graph of the quantification of esterified BODIPY C12 levels is shown. BODIPY C12 levels at each time point were quantified relative to time 0 for each cell line. Data represent mean ± SD of three biological replicates. *\*\*\*\*p<0.0001 by two-way ANOVA with Dunnett’s multiple comparisons test*. D) Representative TLC resolving esterified BODIPY C12 558/568 in Huh7 cells expressing a safe targeting sgRNA or sgRNAs against CLCC1. Cells were incubated with BODIPY C12 for 16 h followed by triacsin C treatment for the indicated times, followed by lipid extraction and TLC. A graph of the quantification of esterified BODIPY C12 levels is shown. BODIPY C12 levels at each time point were quantified relative to time 0 for each cell line. Data represent mean ± SD of three biological replicates. *\*\*\*\*p<0.0001 by two-way ANOVA with Dunnett’s multiple comparisons test*. E) Quantification of ALT from clinical analyzer. Data represent mean ± SD of > four mice. *\*\*\*\*p<0.0001 by one-way ANOVA with Dunnett’s multiple comparisons test*.

**Extended Data Figure 5.**
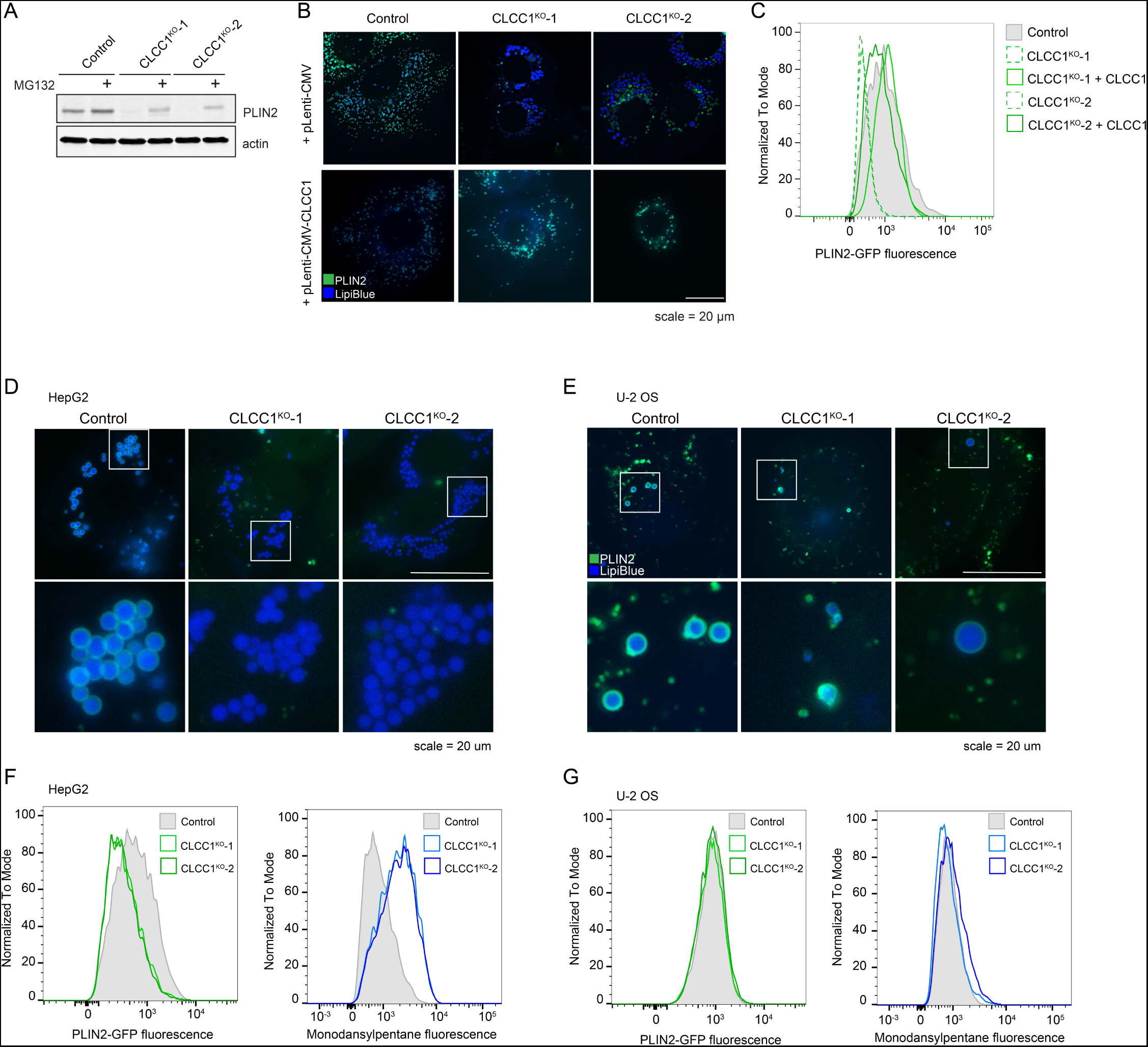
Loss of CLCC1 results in the accumulation of PLIN2-negative lipid droplets. A) Immunoblot of PLIN2-GFP levels in control and CLCC1^KO^ cells treated with 1 µM MG132 for 12 h. B) Representative fluorescence microscopy images of PLIN2-GFP Huh7 cells, control and CLCC1KO cells, expressing a control plasmid or an untagged CLCC1 expression plasmid. LDs are stained with LipiBlue (blue). Scale bar represents 10 µm. C) Flow cytometry histograms of PLIN2-GFP fluorescence in control and CLCC1^KO^ PLIN2-GFP Huh7 cells expressing a control plasmid (dashed) or an untagged CLCC1 expression plasmid (solid). D) Representative fluorescence microscopy images of PLIN2-GFP HepG2 cells expressing a safe targeting sgRNA (control) or sgRNAs against CLCC1. LDs were stained with 500 nM Lipi-Blue. Scale bar represents 20 µm. E) Representative fluorescence microscopy images of PLIN2-GFP U2-OS cells expressing a safe targeting sgRNA (control) or sgRNAs against CLCC1. LDs were stained with 500 nM Lipi-Blue. Scale bar represents 20 µm. F) Flow cytometry histograms of PLIN2 and neutral lipid levels (monodansylpentane fluorescence) in control and CLCC1^KO^ HepG2 cells. G) Flow cytometry histograms of PLIN2 and neutral lipid levels (monodansylpentane fluorescence) in control and CLCC1^KO^ HepG2 cells.

**Extended Data Figure 6.**
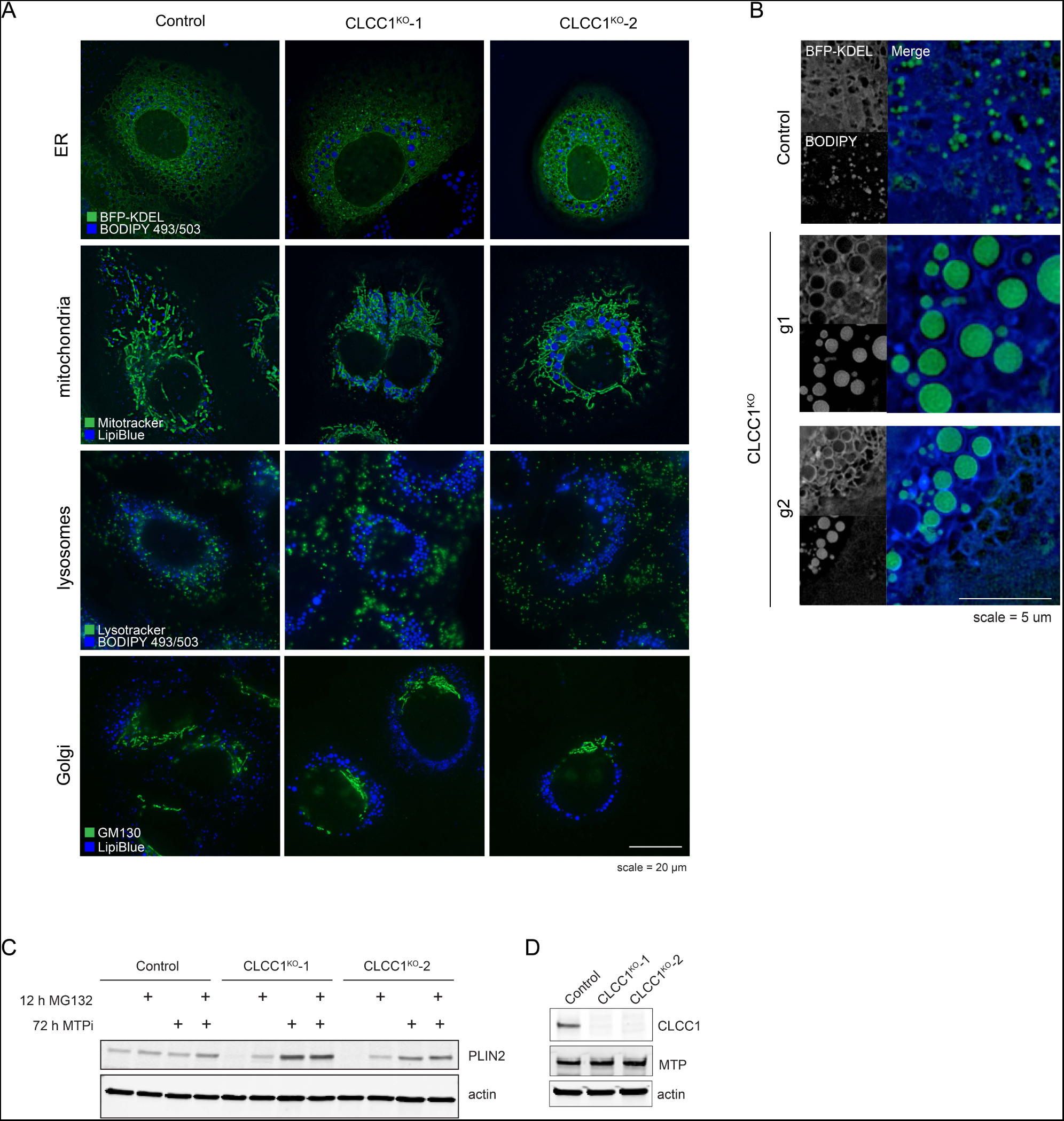
Analysis of organelle morphology in CLCC1 knockout cell lines. A) Representative fluorescence microscopy images of mitochondria, lysosomes, Golgi, and LDs in CLCC1^KO^ cells. Mitochondria was labeled with 500 nM mitotracker green, lysosomes were labeled with 75 nM lysotracker dnd-22, the Golgi apparatus was labeled with rabbit anti-GM130 antibody, and LDs were stained with either 500 nM Lipi-Blue or 1 µg/mL BODIPY 493/503. Scale bar represents 20 µm. B) Fluorescence imaging of ER and LDs in CLCC1^KO^ cells. ER is labeled with a transiently transfected BFP-KDEL construct and LDs were stained with 1 µg/mL BODIPY 493/503. Scale bar represents 5 µm. C) Immunoblot of PLIN2-GFP levels in control and CLCC1^KO^ cells treated with 1 µM MG132 for 12 h and/or 50 nM MTPi for 72 h. D) Immunoblot of MTP levels in control and CLCC1^KO^ cells.

**Extended Data Figure 7.**
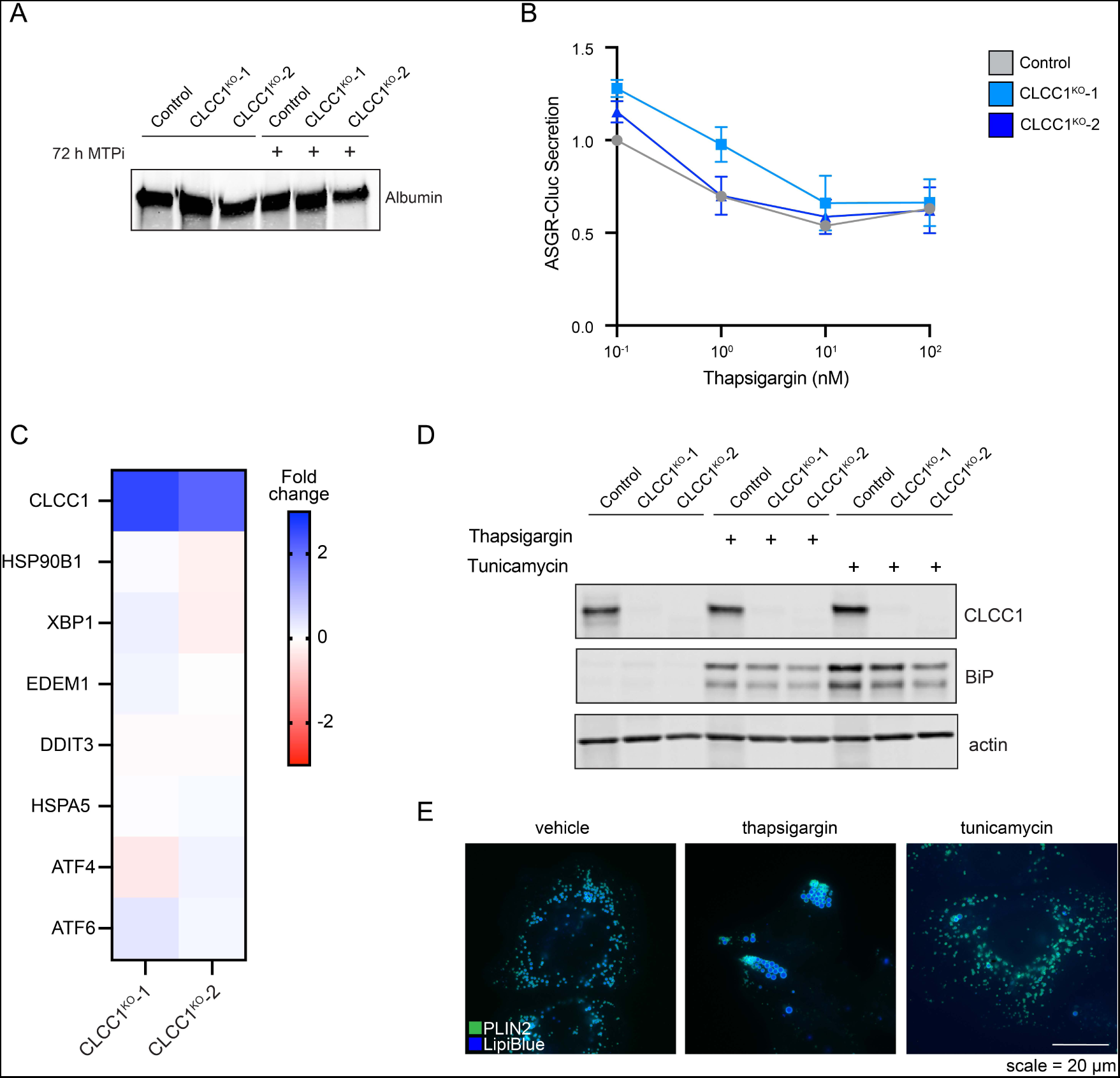
ER stress and general secretion are unchanged in CLCC1 knockout cell lines. A) Immunoblot of albumin secretion from control and CLCC1^KO^ cells. Conditioned serum-free media was collected and precipitated before immunoblotting. B) Quantification of ASGR-Cluc secretion in CLCC1^KO^ cells across three biological replicates. C) Fold change of selected mRNA transcripts in CLCC1^KO^ cells relative to control cells, measured using RNA sequencing. D) Immunoblot of BiP levels in control and CLCC1^KO^ cells treated with 5 µg/mL tunicamycin or 1 µM thapsigargin for 24 h. E) Representative fluorescence microscopy images of PLIN2 (green) and LDs (blue) in Huh7 cells treated with ER stress inducers thapsigargin and tunicamycin. Scale bar represents 20 µm.

**Extended Data Figure 8.**
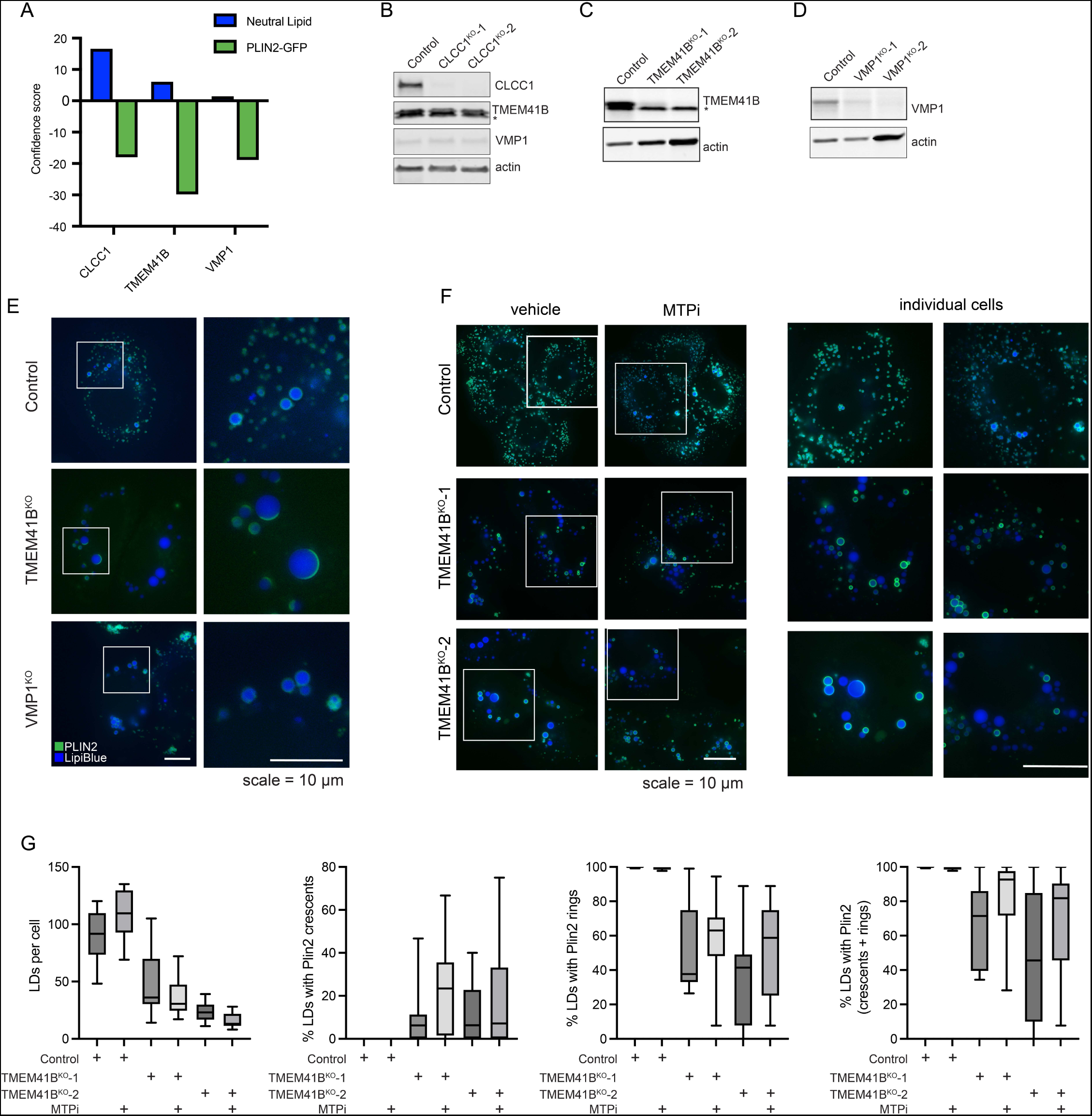
Analysis of ER scramblases and lumenal lipid droplets. A) Confidence score for CLCC1, TMEM41B, and VMP1 from two genome wide CRISPR screens, the neutral lipid (i.e., BODIPY 493/503) screen in the current manuscript and a previous PLIN2-GFP screen^9^. The sign, positive or negative, indicates the effect of gene depletion on neutral lipids or PLIN2-GFP levels. B) Immunoblot of the indicated proteins in control and CLCC1^KO^ Huh7 cells. C) Immunoblot of TMEM41B in control and TMEM41B^KO^ Huh7 cells. D) Immunoblot of VMP1 in control and VMP1^KO^ Huh7 cells. E) Representative fluorescence microscopy images of PLIN2 (green) and LDs (blue) in control and TMEM41B^KO^ and VMP1^KO^ cells. Scale bar represents 10 µm. F) Representative fluorescence microscopy images of PLIN2 (green) and LDs (blue) in control and TMEM41B^KO^ cells treated with MTP inhibitor (MTPi) for 72 h. Zoom images of the boxed regions are shown on the right. Scale bar represents 10 µm. G) Quantification of LDs and LD PLIN2 staining of control and TMEM41B^KO^ cells incubated in the presence and absence of MTP inhibitor (MTPi) as in panel F.

**Extended Data Figure 9.**
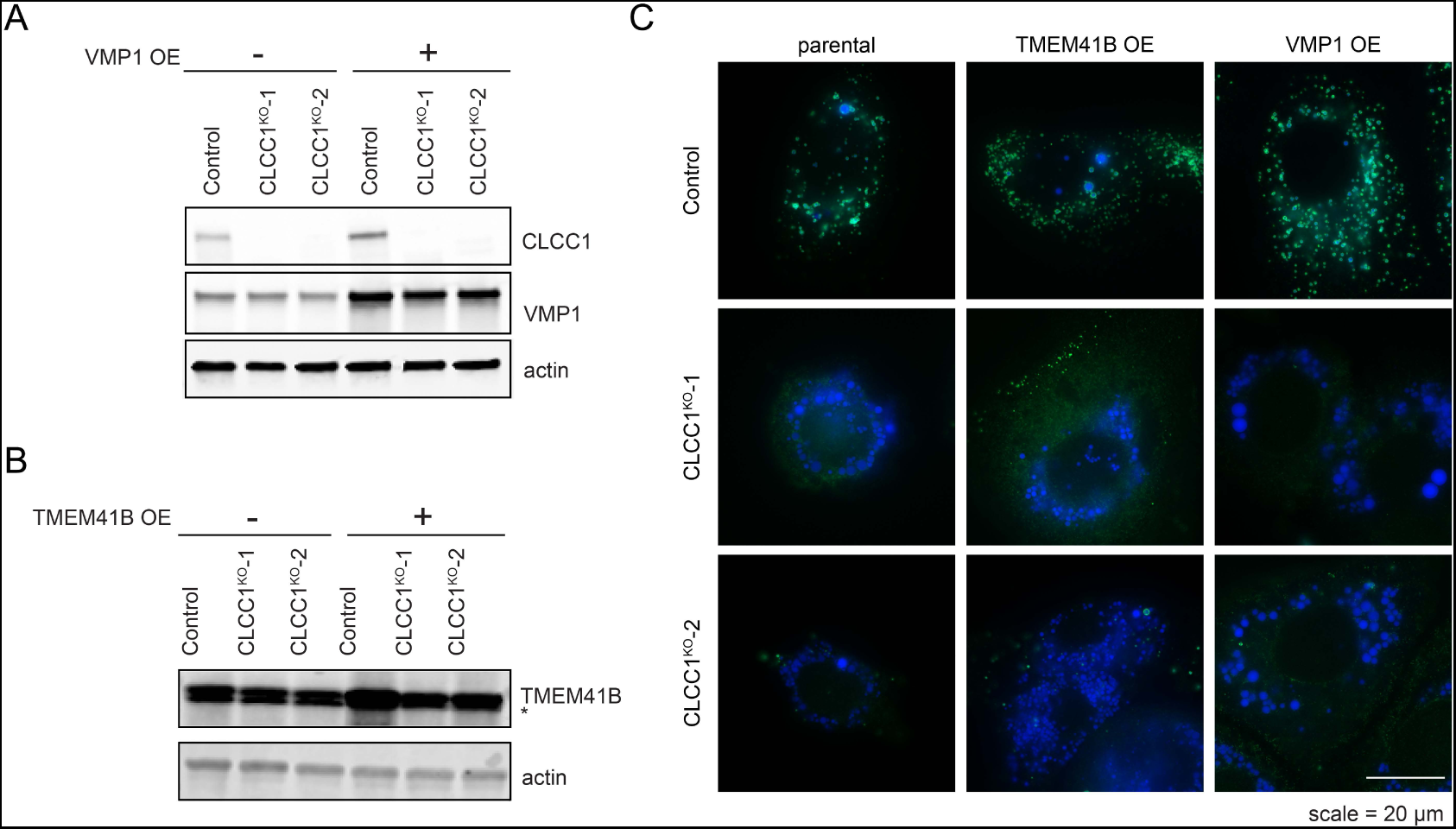
Analysis of ER scramblases in CLCC1^KO^ cells. A) Immunoblot of VMP1 overexpression in control and CLCC1^KO^ cells. B) Immunoblot of TMEM41B overexpression in control and CLCC1^KO^ cells. C) Representative fluorescence microscopy images of PLIN2 (green) and LDs (blue) in control and CLCC1^KO^ cells overexpression TMEM41B and VMP1, as indicated. Scale bar represents 20 µm.

**Extended Data Figure 10.**
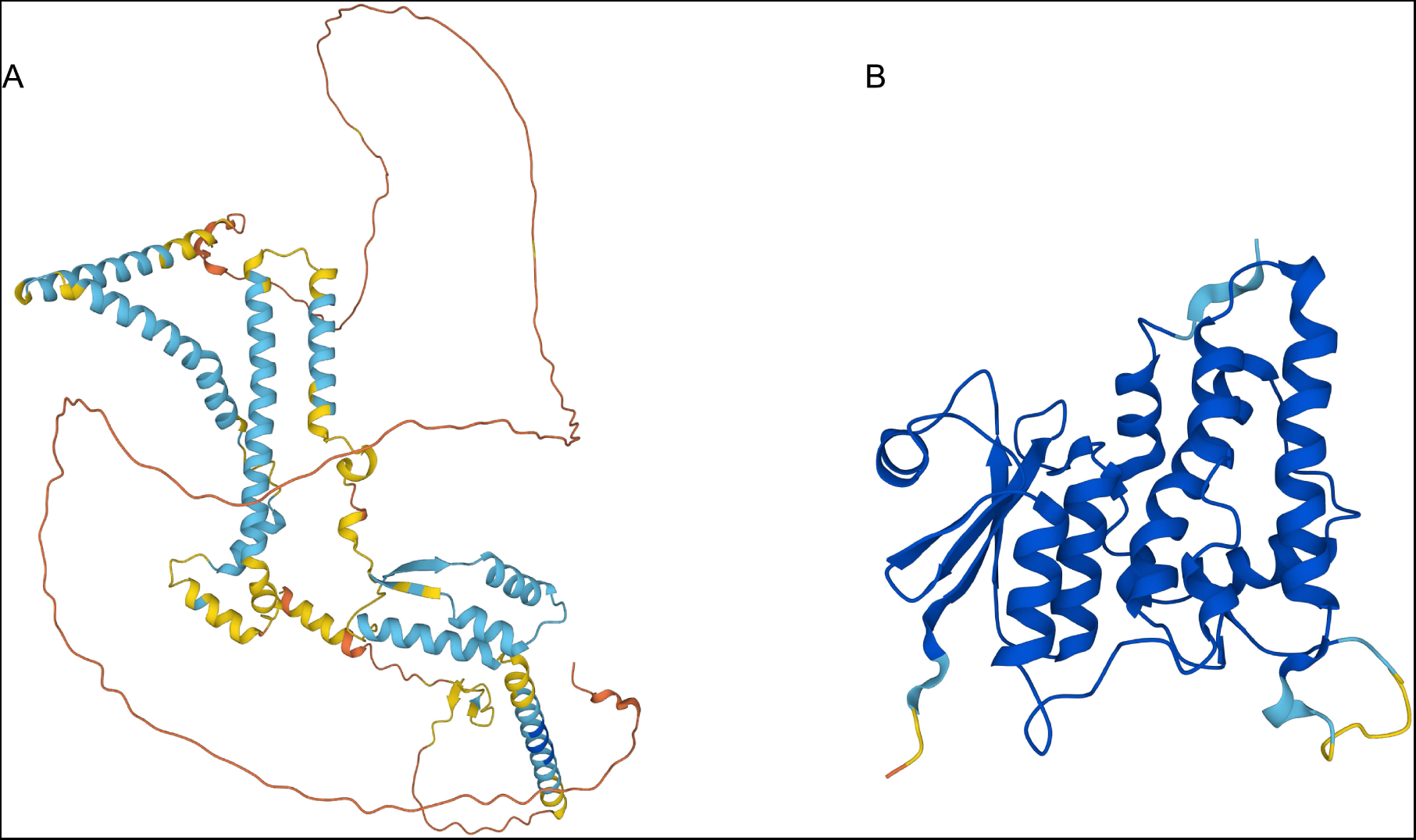
Alphafold and multimer structural predictions. A, B) Alphafold structural predictions of CLCC1 and CLIC1. Images colored by pLDDT (predicted local distance difference test), where blue indicates a confident prediction (light – high, dark – very high), while yellow/orange/red represent predictions of progressively low confidence.

**Extended Data Figure 11.**
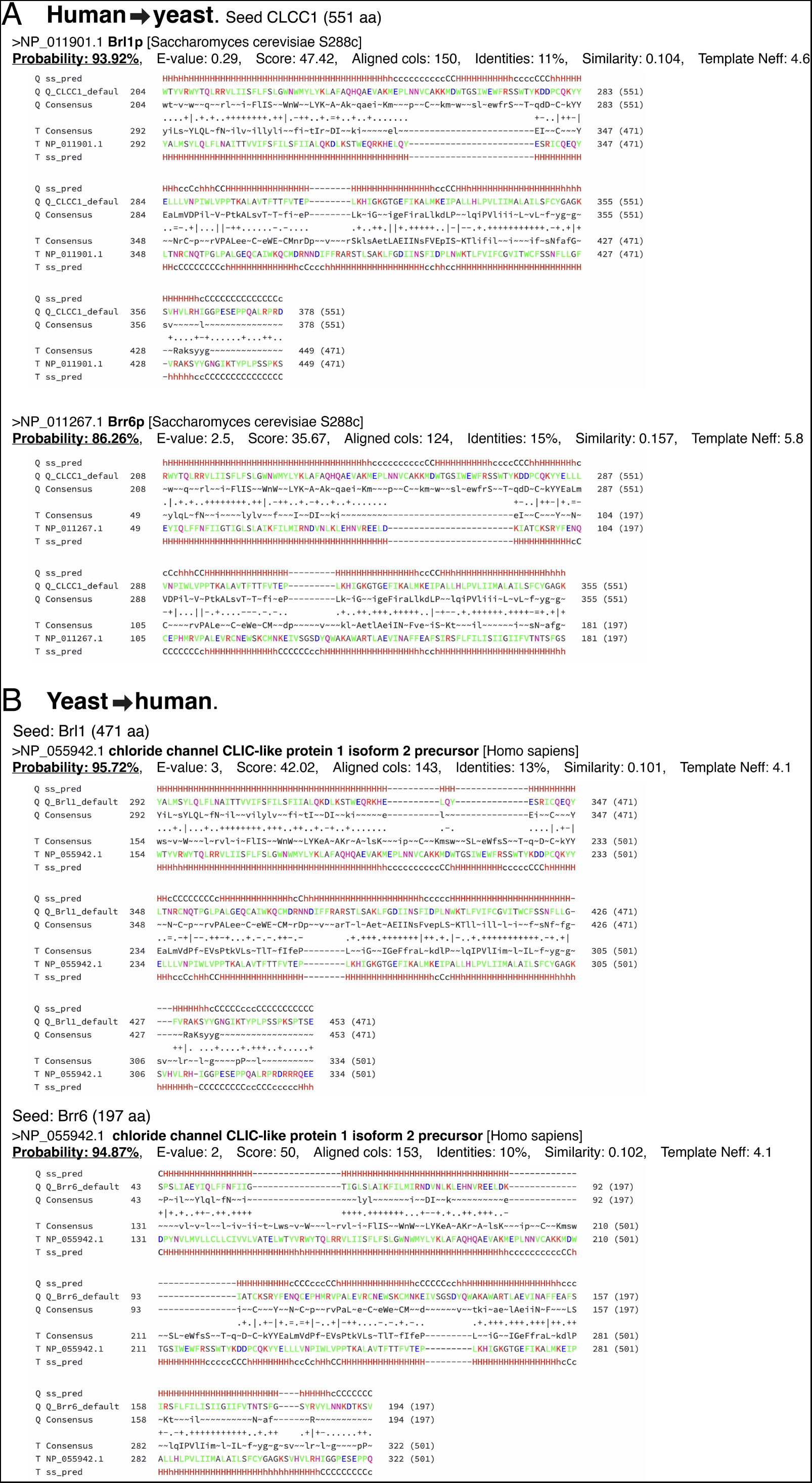
Remote homology analysis linking CLCC1 with Brl1p and Brr6p. HHpred remote homology searches using default settings: (A) in S. cerevisiae using human CLCC1 as the query protein; (B) in human using S. cerevisiae Brl1p or Brr6p as query proteins.

**Extended Data Figure 12.**
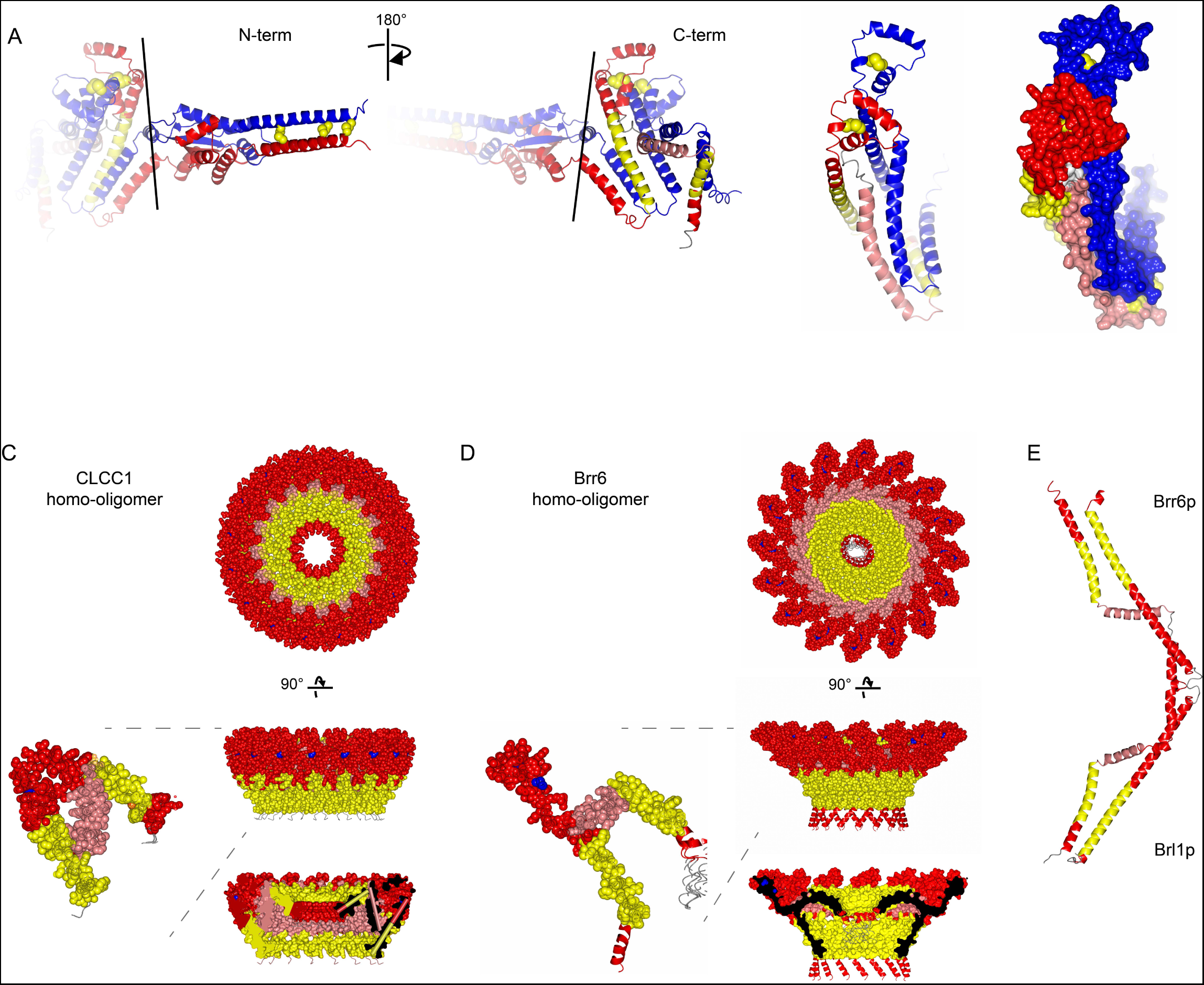
Structure homology analysis of CLCC1 with Brr6p and Brl1p. A) Additional visualizations highlighting the N- and C-termini of the predicted disulfide-stabilized CLCC1 dimer structure Q96S66_V1_5 created by the Levy lab^36^ (downloaded from 3D-Beacons database). The two protomers are colored as in Fig 4C. B) Colabfold structural prediction of CLCC1 (205-360aa) homo-oligomer (16-subunits). Colors: yellow =TMH, pink = AH, red = other helix, blue spheres = conserved cysteines. C) Colabfold structural prediction of Brr6p (28-197aa) homo-oligomer (16-subunits). Colors as B. D) RoseTTAFold structural prediction of Brr6p/Brl1p heterodimer (38-197aa & 281-438)^45^. Colors as B.

**Extended Data Figure 13.**
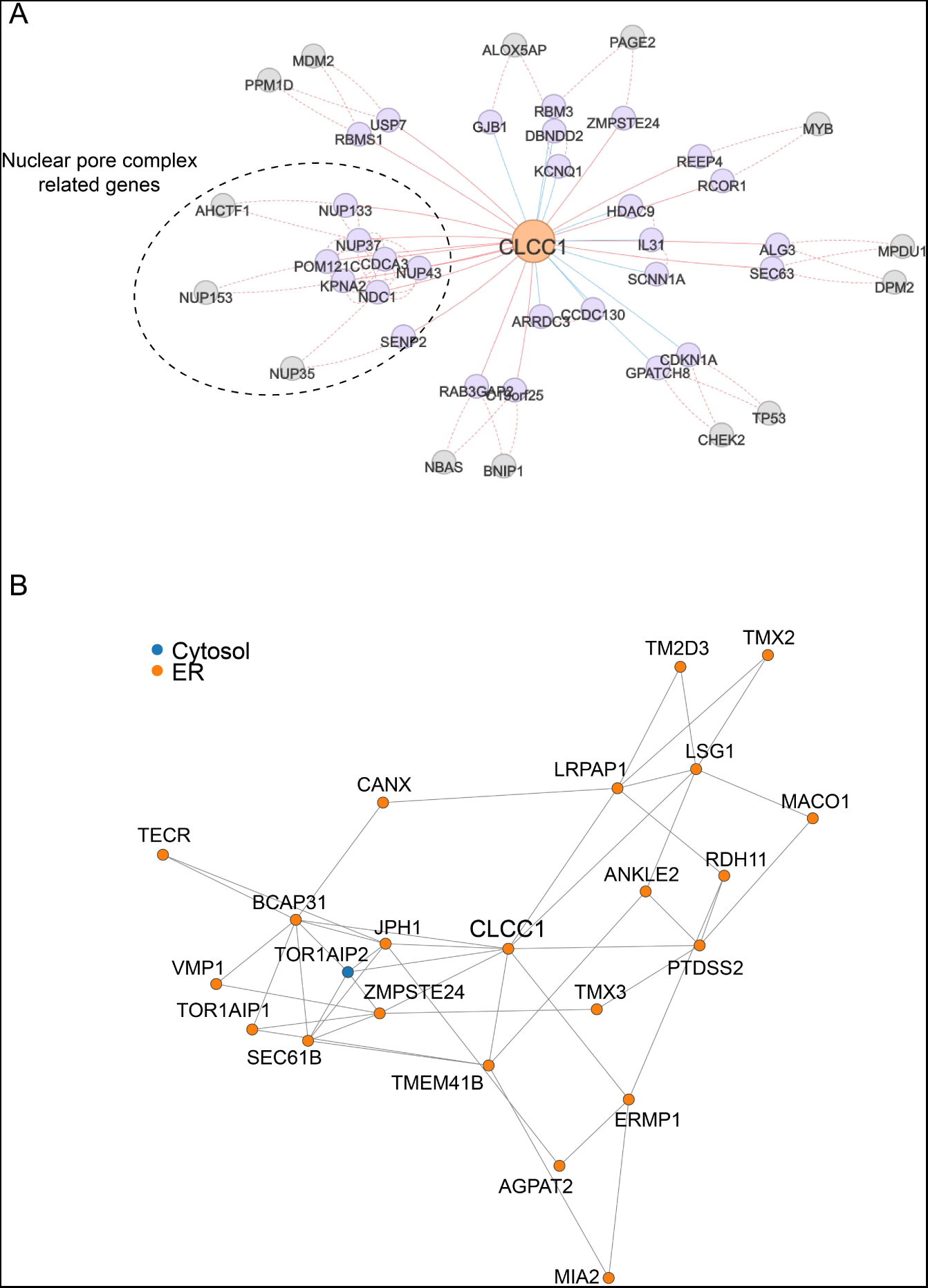
Analyses of CLCC1 co-essentiality and localization neighborhood. A) CLCC1 co-essentiality network using FIREWORKS interactive web tool to reveal gene-gene relationships^41^. B) CLCC1 localization neighborhood using proteomic profiling data of affinity purified organelles^42^.

**Extended Data Figure 14.**
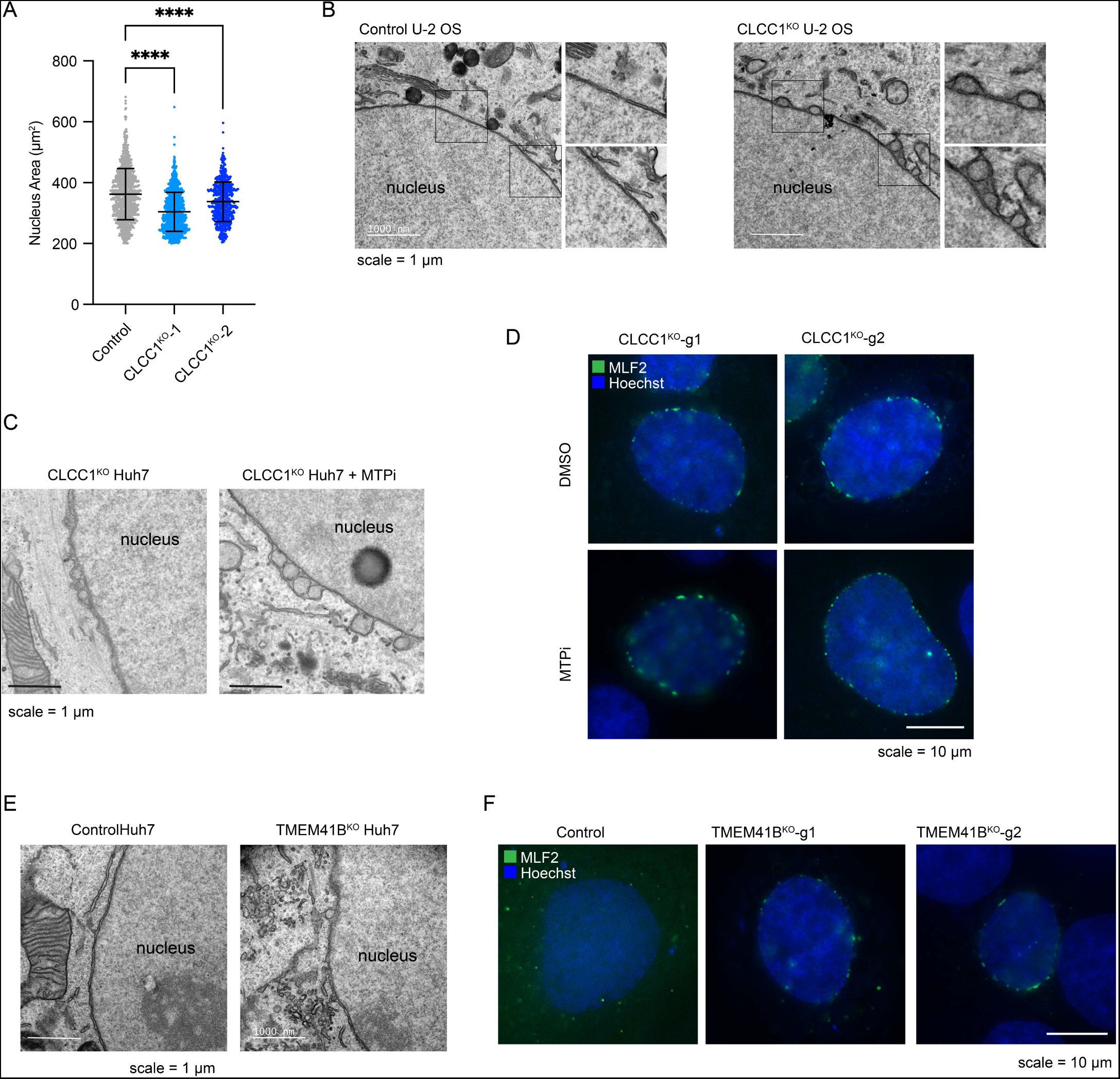
Analysis of nuclear morphology and NE herniations. A) Nuclei were visualized using DAPI fluorescence imaging and the area of the nucleus in control and CLCC1^KO^ cells was quantified. B) Transmission EM of negative stained control and CLCC1^KO^ U-2 OS cells. C) Transmission EM of negative stained CLCC1^KO^ Huh7 cells treated in the presence and absence of MTP inhibitor (MTPi) for 72 h. D) Fluorescence imaging of MLF2-GFP (nuclear bleb marker) in CLCC1^KO^ Huh7 cells treated in the presence and absence of MTP inhibitor (MTPi) for 72 h. E) Transmission EM of negative stained control and TMEM41B^KO^ Huh7 cells. F) Fluorescence imaging of MLF2-GFP (nuclear bleb marker) in control and TMEM41B^KO^ Huh7 cells.

**Extended Data Figure 15.**
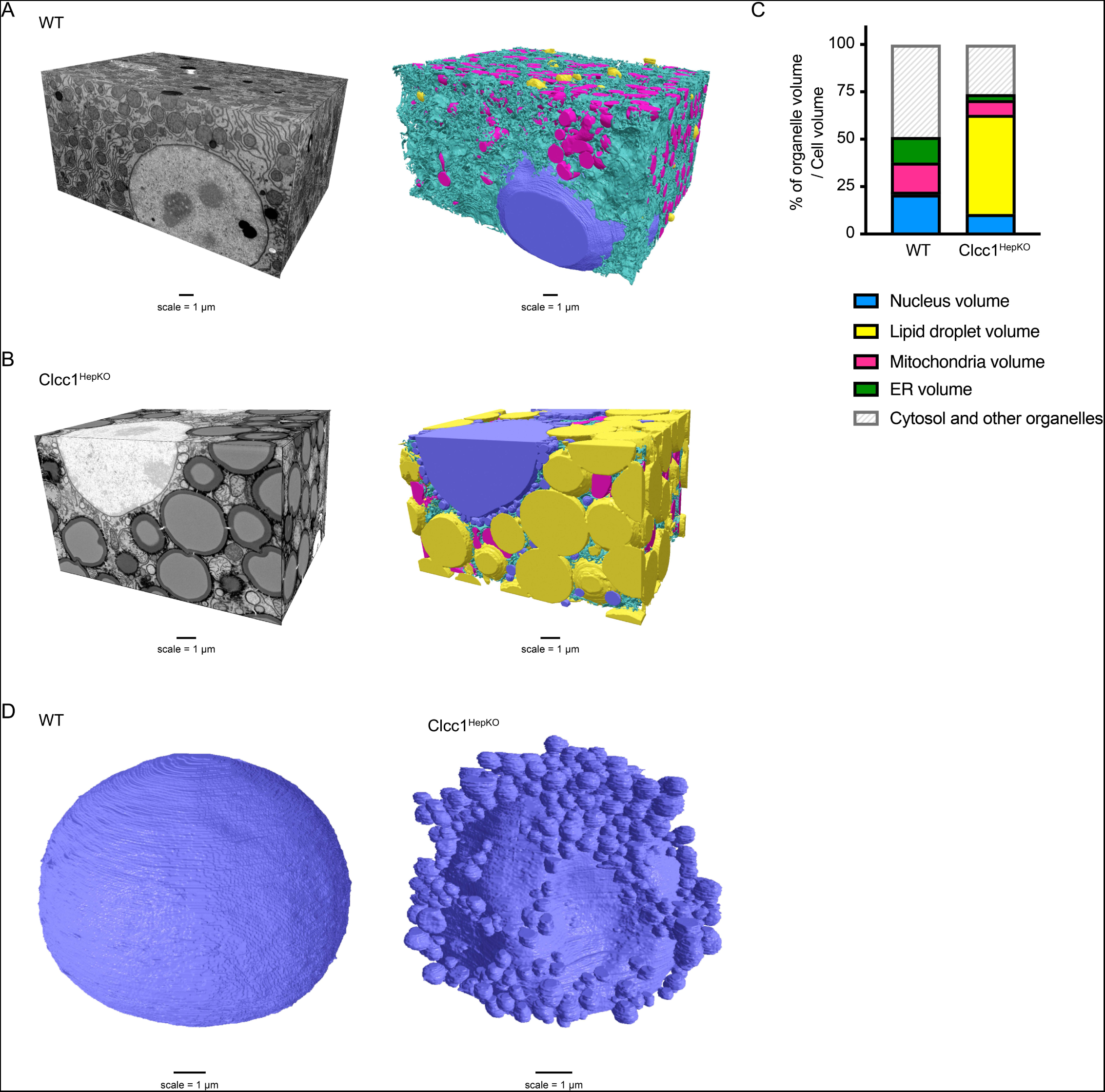
Analysis of organelle architecture using FIB-SEM. A,B) 3D reconstruction of FIB-SEM images and convolutional neural network based automated segmentation of liver volumes derived from liver volumes from WT (A) and Clcc1^HepKO^ (B) mice. ER (Cyan), mitochondria (magenta), LDs (yellow), and nucleus (purple). C) Quantification of organelle volume as a percentage of cellular volume in FIB-SEM reconstructions (A,B) from WT and Clcc1^HepKO^ mice. D) Reconstruction of segmented raw FIB-SEM data for nuclei (blue) from hepatocytes of WT and Clcc1^HepKO^ mice.

**Extended Data Figure 16.**
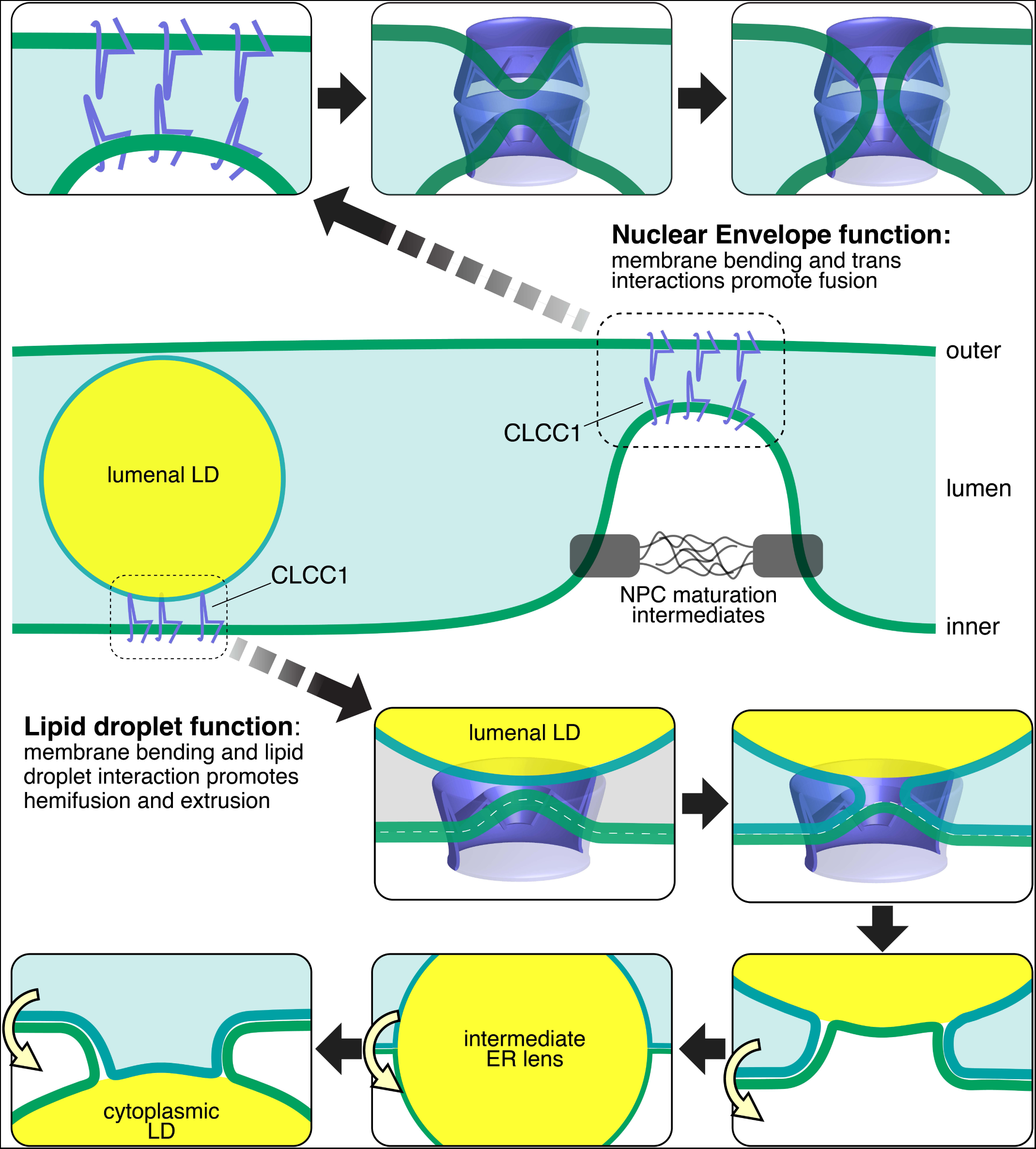
Models of CLCC1 actions at the nuclear envelope and in the ER. Our findings provide strong evidence for a role of CLCC1 in nuclear pore assembly and ER neutral lipid channeling in hepatocytes. Similar to Brl1p and Brr6p in yeast, we propose that CLCC1 acts at the membrane fusion step of nuclear pore complex assembly, which is essential for the insertion of nuclear pores during interphase. Following the insertion of nuclear pore subunits into the inner nuclear membrane, CLCC1 may be recruited to form homo-oligomeric rings that face each other on opposing membranes. Each CLCC1 ring is predicted to bend the membrane towards the other across the lumen, potentially facilitating membrane fusion and perforation. Oligomeric rings formed by Brl1p and its homologs would fulfil the same functions in yeast and organisms in all five eukaryote supergroups^54^. Within the ER, similar structural features of CLCC1 or CLCC1 homo-oligomers promote correct ER neutral lipid flux. In the absence of CLCC1 there is an aberrant increase in neutral lipid flux towards the ER lumen. One possible model is that CLCC1 reduces lumenal LD accumulation by mediating membrane fusion between the inner leaflet of the ER and the lumenal LD. This fusion would form a membrane bridge, allowing neutral lipids to migrate from the lumenal LD into the ER membrane, and emerge in cytosolic LDs. ER scramblases such as TMEM41B could act at this step to ensure that there are sufficient phospholipids for cytosolic emergence. This could represent an LD salvage pathway to provide a homeostatic balance for maintenance of correct amounts of lumenal and cytosolic LDs.

## Supplemental Table Legends

**Supplementary Table 1. CRISPR-Cas9 genetic screen data.**

Full casTLE results for the CRISPR-Cas9 screen employing the genome-wide library of sgRNAs and batch retest sublibrary of sgRNAs under different metabolic conditions.

**Supplementary Table S2. Lipid droplet and metabolism batch retest library.**

List of target genes, sgRNA sequences, and primer sequences used for amplification.

**Supplementary Table S3. Proteomics of LD-enriched buoyant fractions.**

List of proteins identified in proteomics analyses of LD-enriched buoyant fractions isolated from control and CLCC1^KO^ cells.

